# Hippo-activated cells induce non-cell autonomous tumorigenesis in *Drosophila*

**DOI:** 10.1101/2025.05.05.652237

**Authors:** Daichi Honda, Misako Okumura, Chisako Sakuma, Toshinori Ando, Masayuki Miura, Takahiro Chihara

## Abstract

The Hippo pathway is known as a tumor-suppressor pathway, and most related studies have indicated that the inhibition of the Hippo pathway leads to tumorigenesis. However, recent studies have suggested that the activated Hippo pathway can potentially promote tumorigenesis in some contexts. Here, we show that the activated Hippo pathway induces non-cell-autonomous tumorigenesis characterized by tumor markers in the *Drosophila* wing epithelium. This suggests that the Hippo-activated cells behave similarly to “oncogenic niche cells.” We found that the Hippo-activated cells distantly induced tumor-like cells exhibiting aberrant cell proliferation. Moreover, we identified the evolutionarily conserved amino acid transporters, Sat1/2, that redundantly work with the growth factors, Wingless and Spitz, for non-cell autonomous tumorigenesis. Our findings provide novel insights into the Hippo pathway and may be useful in developing cancer treatments targeting the Hippo pathway.

## Introduction

The Hippo pathway, a highly conserved tumor-suppressor pathway, negatively regulates organ growth and tumorigenesis (Moroishi, Hansen, and Guan 2015; Yu, Zhao, and Guan 2015). The *Drosophila* Hippo pathway consists of the main components, Hippo (Hpo) and Warts (Wts), which target the transcriptional coactivator Yorkie (Yki: Yes-associated protein, YAP in mammals) (Moroishi, Hansen, and Guan 2015; Yu, Zhao, and Guan 2015). Given that the activated Hippo pathway suppresses Yki by sequestrating and degrading Yki in the cytosol, the inactivation of the Hippo pathway allows Yki to translocate into the nucleus and initiate tumorigenesis by promoting cell proliferation, cell competition, and migration (Ziosi et al. 2010; Thompson and Cohen 2006; Ding et al. 2021; Nagata et al. 2022; Ding et al. 2021; Udan et al. 2003).

In contrast to this classic model in which the Hippo pathway acts as a tumor suppressor, recent reports suggest that the Hippo pathway acts as a tumor promoter. For instance, LATS1/2 (human orthologs of Wts), which are those of the Hippo pathway components, suppresses anti-tumor responses by inhibiting the secretion of signal molecules to activate the immune cells in murine cancer cells (Moroishi et al. 2016). Interestingly, LATS1/2 behaves as a tumor promoter *in vivo* but not *in vitro* (Moroishi et al. 2016), calling into question the role of the Hippo pathway in the actual body. Furthermore, YAP is frequently silenced in multiple cancers (Carter et al. 1994; Hampton et al. 1994; Cottini et al. 2014; Yuan et al. 2008; Pearson et al. 2021). For instance, YAP loss promotes tumor growth in breast cancers and migration in hematological cancers (Carter et al. 1994; Hampton et al. 1994; Cottini et al. 2014; Yuan et al. 2008). These recent studies suggest that, in some contexts, Hippo activation/YAP silencing functions as a tumor promoter. However, the mechanism underlying the tumorigenesis caused by Hippo activation remains poorly understood.

Here, we investigated the tumor-promoting role of the Hippo pathway using the *Drosophila* wing disc, where tumorigenesis can be analyzed by genetic manipulations (Martínez-Abarca Millán et al. 2023; Gandille et al. 2010). We found that Hippo activation can induce tumors characterized by phospho-S6 staining (tissue growth), matrix metalloproteinase 1 (MMP1) expression (invasive ability), and 5-ethynyl-2′-deoxyuridine (EdU) staining (excessive cell proliferation) in a non-cell-autonomous manner. This tumorigenesis occurs through the Hippo target genes, including *rpr*, *hid, grim* (an apoptotic inducer), *atg8a* (an autophagy-related gene), *cyclin E* (a proliferative activator), and microRNA *bantam*. We found that the signaling pathway of the compensatory proliferation, Dronc-Wg/Spitz signaling, acting downstream of Rpr, Hid, and Grim is required for Hippo activation-mediated tumorigenesis. Moreover, we identified that the evolutionarily conserved amino acid transporters, Strip-interacting amino acid transporters 1/2 (Sat1/2), were required and redundantly worked with growth factors for Hippo activation-mediated tumorigenesis. Therefore, we suggest that the Hippo-activated cells cause non-cell-autonomous tumorigenesis through the growth factors Wg/Spitz and Sat1/2.

## Result

### The activated Hippo pathway induces tumorigenesis

First, we genetically induced the Hippo-activated cells in the *Drosophila* epithelial tissue, wing disc, using the *ptc-Gal4* driver that is expressed in the anterior/posterior (A/P) compartment boundary. Given that the Hippo pathway is suppressed by the striatin-interacting phosphatase and kinase (STRIPAK) complex, the knockdown of a STRIPAK component *strip* can effectively activate the Hippo pathway (Chen et al. 2019, Fig. 1A). Additionally, *wts* overexpression or *yki* knockdown can induce the Hippo-activated phenotypes (Huang et al. 2005; Oh and Irvine 2008, Fig. 1A). We examined the effects of Hippo activation on tumorigenesis using the following markers: the activation of the mTOR pathway, MMP1 expression, and EdU uptake. It is known that the mTOR signaling, MMP1 expression, and EdU uptake for tissue growth, invasive ability, and cell proliferation are upregulated in the tumorigenesis process of *Drosophila* and mammals (Nagata et al. 2022; Zhang et al. 2022; Uhlirova and Bohmann 2006; Zou et al. 2020; Parker and Struhl 2020).

**Figure 1.**
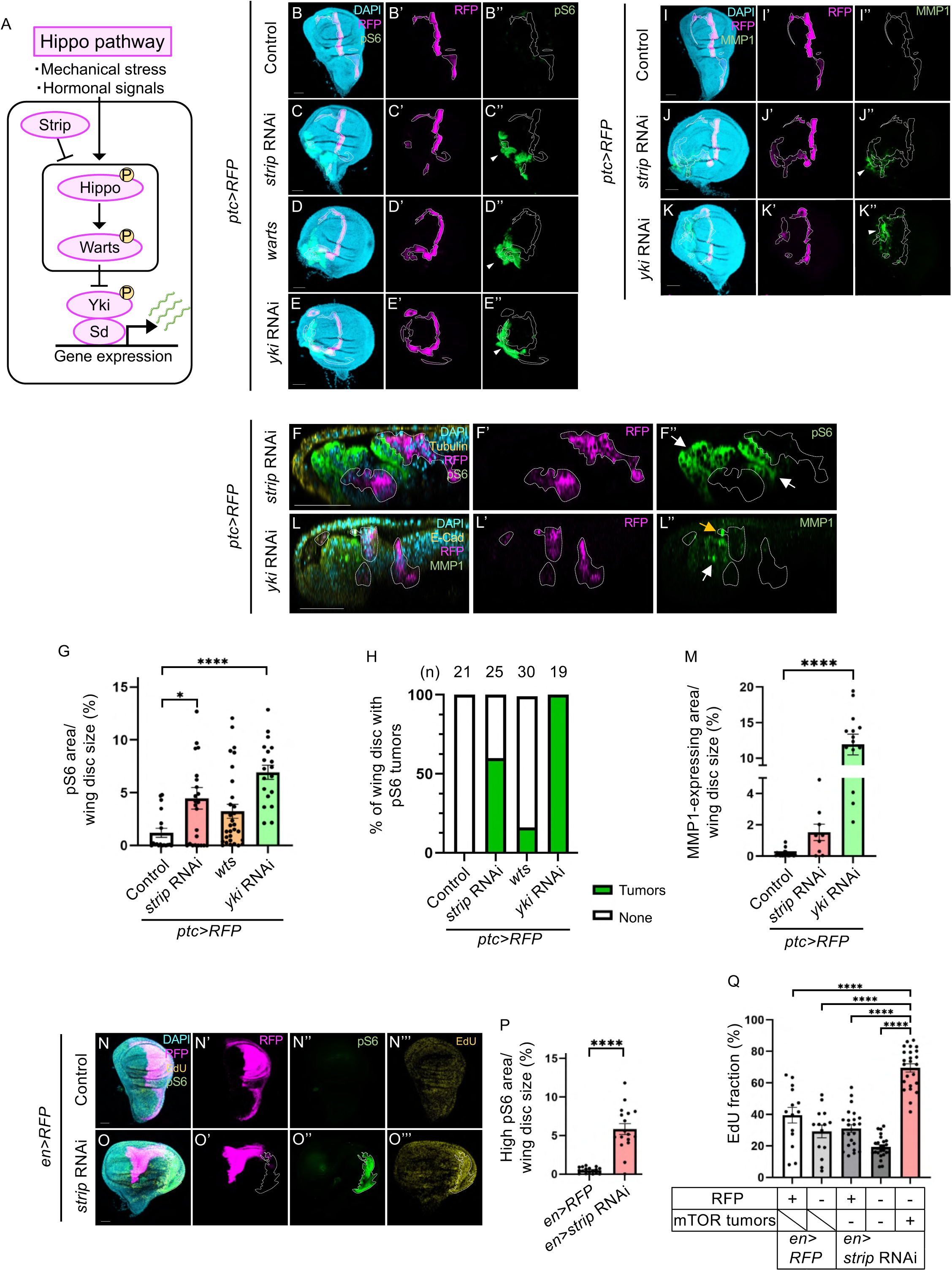
The activated Hippo pathway upregulates the mTOR pathway, MMP1 expression, and EdU uptake in a non-cell-autonomous manner. (A) The schematic depiction indicates the Hippo pathway and its function. (B–E) Confocal images show wing discs bearing wild-type, *strip-*, *yki-*knockdown, or *wts*-overexpressing cells marked with RFP expression (magenta) and stained with anti-phospho-S6 (green). The white arrowhead shows the mTOR-activated tumors. (F) Confocal images show wing discs bearing *strip-*knockdown cells marked with RFP expression (magenta) and stained with anti-phospho-S6 (green) and anti-tubulin (yellow). The XZ section images of the wing disc in Supplemented Fig. 1B. The white arrow indicates non-cell-autonomous activation of the mTOR pathway. RFP (magenta) outlined by white dashed lines marks the expression pattern of *ptc-Gal4* in the wing disc (B–F). (G) Quantification of the size of the phospho-S6 positive region (% of phospho-S6 positive area/disc area) in wild-type, *strip-, yki*-knocked down, or *wts-*overexpressed wing disc. ∗∗∗∗p < 0.0001; ∗p < 0.05; one-way ANOVA with Dunnett’s multiple comparison test. (H) Incidence of mTOR (phospho-S6)-activated tumors. n = 21 (*ptc > RFP*), n = 25 (*ptc > strip* RNAi), n = 30 (*ptc > wts*), n = 19 (*ptc > yki* RNAi). (I–K) Confocal images show the wing discs bearing wild-type, *strip-*, or *yki-*knocked down cells marked with RFP expression (magenta) and stained with anti-MMP1 (green). The white arrowhead shows the MMP1-expressing cells. (L) Confocal images show wing discs bearing *strip-*knockdown cells marked with RFP expression (magenta) and stained with anti-MMP1 (green) and anti-E-Cad (yellow). The XZ section images of the wing disc in Supplemented Fig. 1D. The white arrow indicates non-cell-autonomous MMP1 expression. The yellow arrow indicates cell-autonomous MMP1 expression. RFP (magenta) outlined by white dashed lines marks the expression pattern of *ptc-Gal4* in the wing disc (I–K, L). (M) Quantification of the MMP1-expressing area (% of MMP1-expressing area/disc area) in wild-type, *strip*-, or *yki*-knocked down wing disc. ∗∗∗∗p < 0.0001; one-way ANOVA with Dunnett’s multiple comparison test. (N–O) Confocal images show the wing discs bearing wild-type or *strip-*knocked down cells marked with RFP expression (magenta) and stained with anti-phospho-S6 (green) and EdU (yellow). RFP (magenta) marks the expression pattern of *en-Gal4* in the wing disc. mTOR-activated tumors are outlined by white dashed lines. (P) Quantification of the size of the phospho-S6 positive region (% of phospho-S6 positive area/disc area) in wild-type or *strip-*knocked down wing disc. ∗∗∗∗p < 0.0001; Welch’s t-test. (Q) Quantification of the EdU-stained area (% of EdU-stained area/RFP-positive, RFP-negative, or mTOR tumor area) in the wild-type wing disc. ∗∗∗∗p < 0.0001; one-way ANOVA with Dunnett’s multiple comparison test. Scale bars represent 50 μm.

In the wild-type wing disc, mTOR signaling was sparsely and weakly activated, as indicated by the scattered staining with the antibody against the phosphorylation of ribosomal protein S6 (pS6) (Fig. 1B). Contrarily, the Hippo-activated cells induced the strong activation of mTOR signaling (pS6 staining in Fig. 1B–E, quantified in G and H). All three types of Hippo-activated/Yki-suppressed cells, which are the cells of *strip* RNAi, *yki* RNAi, or *wts* overexpression, activated mTOR signaling, although the efficiency of mTOR activation/tumors was different among the three types (Fig. 1G and H). Interestingly, the Hippo-activated cells rarely overlapped with the cells with mTOR activation, and mTOR activation was induced in the around the Hippo-activated cells, indicating that the Hippo signaling activation induces mTOR signaling non-cell-autonomously (Fig. 1F, Supplementary Fig. 1A, B). In addition to mTOR activation, the Hippo-activated cells induced MMP1 expression (Fig. 1I–K, quantified in M). The MMP1 upregulation tended to occur in both *strip*/*yki* RNAi, although the MMP1 upregulation by *yki* RNAi was stronger than that by *strip* RNAi (Fig. 1M). MMP1 upregulation tended to be induced by Hippo activation in a cell-autonomous and non-cell-autonomous manner (Fig. 1L, Supplementary Fig. 1C, D). The MMP1 expression was frequently observed in the mTOR-activated cells in a non-cell-autonomous manner (Supplementary Fig. 1E–L). In addition to *ptc-Gal4*, the mTOR-activated tumors emerged when the Hippo pathway was activated using the *en-Gal4* driver expressed in the posterior compartment (Fig. 1N and O, quantified in P). In the mTOR-activated cells, cell proliferation was strongly upregulated, as indicated by EdU staining (Fig. 1N and O, quantified in Q). These data suggest that the Hippo-activated cells behave like “oncogenic niche cells” that induce non-cell-autonomous tumorigenesis.

### Hippo activation in the hinge/ventral notum regions induces the mTOR-activated tumors

We generated the Hippo-activated cells in the wing disc using *ptc-Gal4*, *en-Gal4*, or *ci-Gal4* drivers, which are expressed in the A/P compartment boundary, anterior compartment, and posterior compartment, respectively. The mTOR-activated tumors tended to emerge in the putative hinge and notum region but not in the pouch region (Fig. 1B–E, N, O, Supplementary Fig. 2A–C), although *strip* RNAi activated the Hippo pathway in not only the hinge but also in the pouch region, as indicated by the suppression of Expanded, a *yki*-target gene (Supplementary Fig. 3A–E, quantified in F). This suggests that there might be a cell-type/region specificity for the non-cell-autonomous tumorigenesis. To examine this possibility, we activated the Hippo pathway in the specific region of the wing disc, including the pouch, hinge, or notum region, by using *nub-Gal4*, *bx-Gal4*, *pnr-Gal4*, *284-Gal4*, or *41D11-Gal4* (the expression pattern of each *Gal4* line is summarized in Fig. 2A). The Hippo activation in the pouch or dorsal notum region by using *nub-Gal4*, *bx-Gal4*, *pnr-Gal4*, or *284-Gal4* did not induce the mTOR-activated tumors (Fig. 2B–D, F, G, quantified in E, H, Supplementary Fig. 2D, E, G, H, quantified in F, I), although apoptosis and attenuation of tissue growth, known as a characteristic of Hippo activation, were observed in the pouch region (Supplementary Fig. 3G, H,, quantified in I, J). Contrarily, when the Hippo pathway was activated in the dorsal hinge and ventral notum region using *41D11-Gal4*, mTOR-activated tumors were observed (Fig. 2I–K, quantified in L). Additionally, we used an *Ay-Gal4* driver (*actin promoter-FRT-y^+^-STOP-FRT-Gal4*) that can randomly generate the Hippo-activated clones in the wing disc (Fig. 2M and N). When the Hippo-activated clones were induced in the ventral/lateral hinge region, the mTOR-activated tumors emerged in those nearby areas (Fig. 2M and N). Therefore, we conclude that the hinge/ventral notum cells can become oncogenic niche cells when the Hippo pathway is activated.

**Figure 2.**
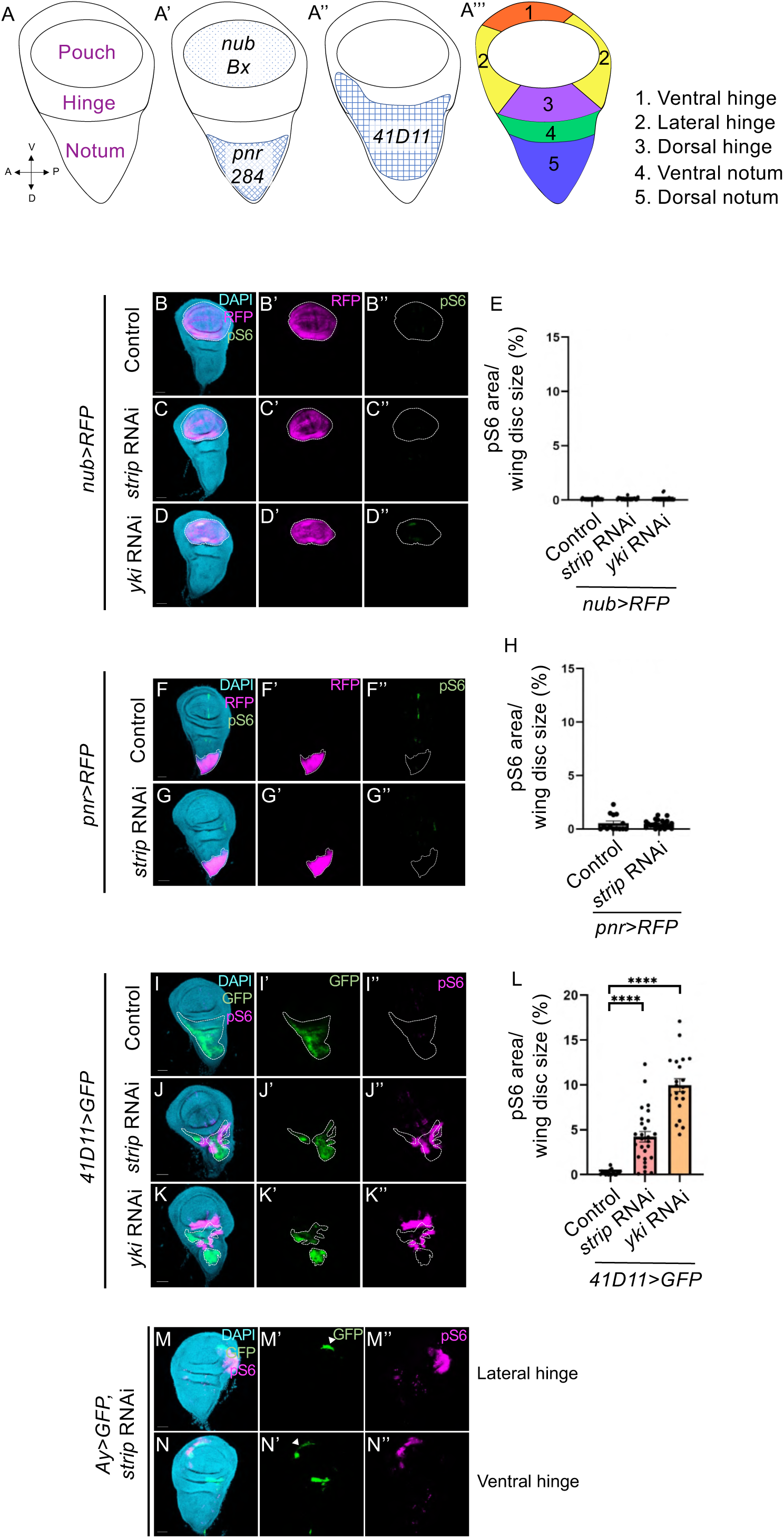
Hippo activation in the hinge/ventral notum region induces mTOR-activated tumors. (A) The schematic depictions indicate the compartments (A), the active area of *nub-Gal4*, *bx-Gal4*, *284-Gal4*, *pnr-Gal4* (A’), and *41D11-Gal4* (A’’), and subcompartment (A’’’) in the wing disc. (B–D, F, G) The confocal images show the wing discs bearing wild-type, *strip-*, or *yki*-knockdown cells marked with RFP expression (magenta) and stained with anti-phospho-S6 (green). RFP (magenta) outlined by white dashed lines marks the expression pattern of *nub-Gal4* or *pnr-Gal4* in the wing disc. (I–K) Confocal images show the wing discs bearing wild-type, *strip-*, or *yki*-knockdown cells marked with GFP expression (green) and stained with anti-phospho-S6 (magenta). GFP (green) outlined by white dashed lines marks the expression pattern of *41D11-Gal4* in the wing disc. (E, H, L) Quantification of the size of the phospho-S6 positive region (% of phospho-S6 positive area/disc area) in the wild-type, *strip-*, or *yki*-knocked down wing disc. Welch’s t-test for (H). ∗∗∗∗p < 0.0001; one-way ANOVA with Dunnett’s multiple comparison test for (E and L). (M–N) The confocal images show wing discs bearing *strip-*knocked down clones marked with GFP expression (green) and stained with anti-phospho-S6 (magenta). GFP (green) marks the expression pattern of *Ay-Gal4* in the wing disc. The white arrowhead indicates the *strip-*knocked down clones in the lateral/ventral hinge. Scale bars represent 50 μm.

### rpr, hid, grim, bantam, cyclin E, and atg8a are involved in non-cell-autonomous tumorigenesis

We investigated how the activated Hippo pathway induces non-cell-autonomous tumorigenesis. Given that the Hippo pathway regulates the expression of multiple genes (Udan et al. 2003; Ziosi et al. 2010; Thompson and Cohen 2006), we investigated which target genes are crucial for Hippo activation-mediated tumorigenesis. To this end, we conducted the overexpression or knockdown of previously reported Hippo/Yki-target genes (Fig. 3A, Udan et al. 2003; Ziosi et al. 2010; Thompson and Cohen 2006; Kowalczyk et al. 2022; D. Wang et al. 2020; Seo et al. 2023). We found that either the *bantam* overexpression, *cyclin E* overexpression, *atg8a* knockdown, or knockdown of *rpr*, *hid*, and *grim* suppressed the non-cell-autonomous mTOR activation caused by Hippo activation (Fig. 3B–I, quantified in J, K). Contrarily, the expression levels of the other Hippo-target genes, *myc, brk*, *chinmo*, *fng*, *diap1*, and *InR,* were not involved in non-cell-autonomous mTOR activation (Supplementary Fig. 4A–G, quantified in H, Fig. 1J). These results indicate that Cyclin E (a proliferative activator), Atg8a (an autophagy-related gene), Rpr, Hid, and Grim (apoptosis inducers), and *bantam* (a regulator of cell proliferation, autophagy, and apoptosis) (Udan et al. 2003; Ziosi et al. 2010; Thompson and Cohen 2006) are involved in the non-cell-autonomous tumorigenesis induced by the Hippo-activated cells.

**Figure 3.**
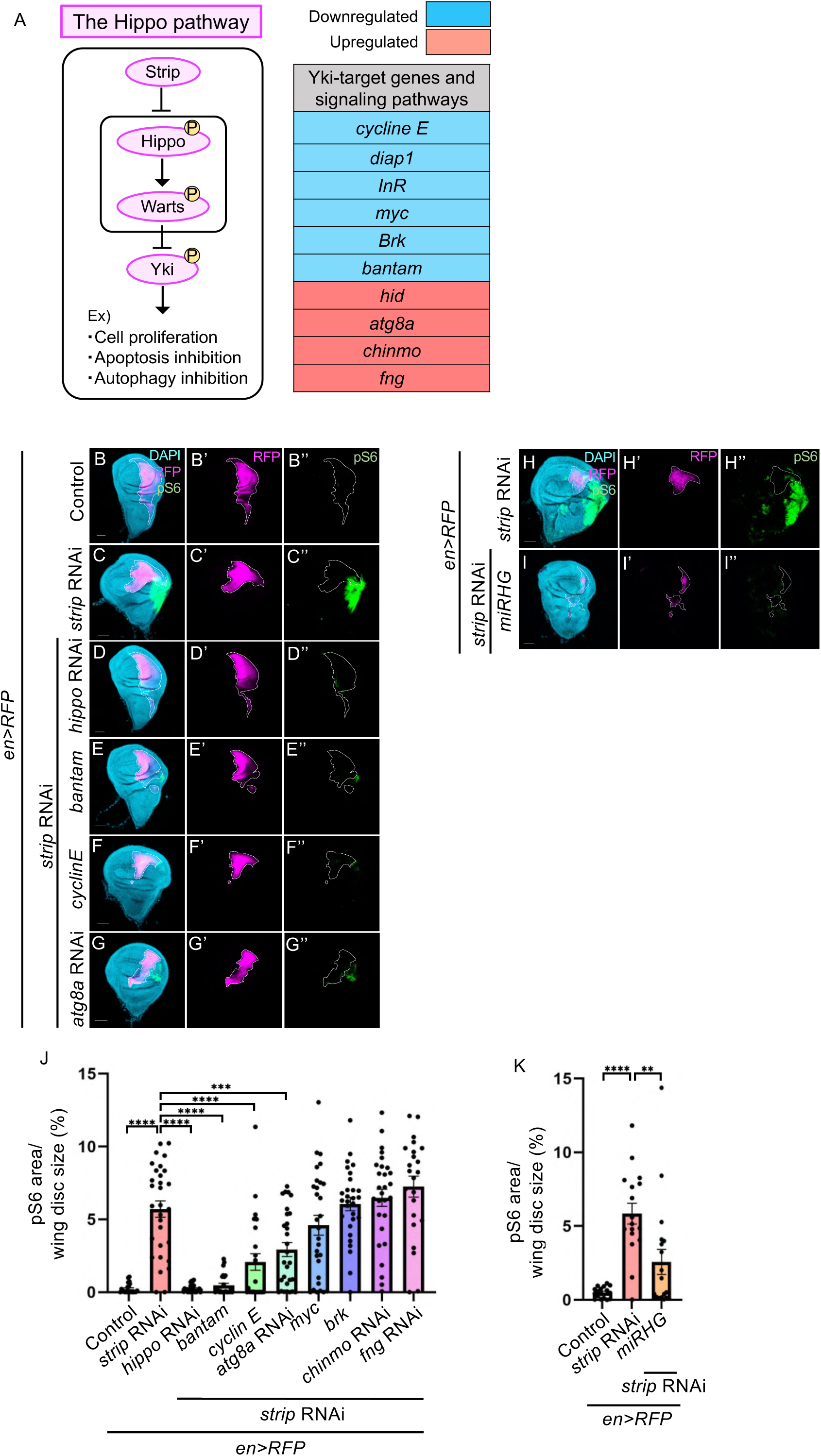
Hippo activation induces non-cell-autonomous mTOR activation by regulating several Hippo-target genes. (A) The left schematic depiction indicates the Hippo pathway and its function. The right table shows the Hippo/Yki-target genes and signaling pathways. The blue- and red-colored genes are downregulated and upregulated when the Hippo pathway is activated. (B–I) The confocal images show the wing discs bearing wild-type, *strip-*knockdown cells with or without knockdown of *hippo, atg8a*, *rpr, hid, grim*, overexpression of *bantam*, or *cyclin E* marked with RFP expression (magenta) and stained with anti-phospho-S6 (green). RFP (magenta) outlined by white dashed lines marks the expression pattern of *en-Gal4* in the wing disc. Knockdown of *rpr, hid,* and *grim* was conducted by the expression of microRNAs for *rpr*, *hid*, and *grim* (*miRHG*). (J–K) Quantification of the phospho-S6 positive region size (% of phospho-S6 positive area/disc area) in the wing disc-bearing wild-type, *strip-*knockdown cells with or without the knockdown of *hippo, atg8a, rpr, hid, grim*, *chinmo*, and *fng*, or overexpression of *bantam*, *cyclin E*, *myc*, or *brk*. ∗∗∗∗p < 0.0001; ∗∗∗p < 0.001; one-way ANOVA with Dunnett’s multiple comparison test. Scale bars represent 50 μm.

### Hippo activation induces non-cell-autonomous tumorigenesis through Dronc-Wg/Spitz signaling

Given that the Hippo pathway drives apoptosis signaling (Udan et al. 2003, Supplementary Fig. 5A), which is also crucial for Hippo activation-mediated tumorigenesis (Fig. 3H, I, quantified in K), we examined how apoptosis signaling is involved in Hippo activation-mediated tumorigenesis. The Hippo pathway promotes *hid* expression that forms a complex with Rpr and Grim to activate the initiator caspase Dronc, which interacts with the caspase activator Dark, and subsequently activates the effector caspase Drice/Dcp-1 for apoptosis (Tian, Li, and Shi 2024; Xu et al. 2006; Udan et al. 2003, Supplementary Fig. 5A). We found that Hippo activation by *strip* RNAi drives apoptosis signaling through Rpr, Hid, and Grim (Supplementary Fig. 5B–D, quantified in F) as well as the previous study conducted by Hippo overexpression (Udan et al. 2003). Next, we investigated the involvement of Dark, Dronc, and Drice/Dcp1 as downstream components of Rpr, Hid, and Grim. The knockdown of *dark* or *dronc* significantly suppressed the non-cell-autonomous mTOR activation (Fig. 4A–C, quantified in F). Contrarily, the expression of *p35*, a Drice/Dcp1 inhibitor, could not suppress, but rather promote, mTOR activation (Fig. 4D, quantified in G), although the *p35* expression could significantly suppress the activated Drice/Dcp1 level (Supplementary Fig. 5E, quantified in F). Therefore, we expect that the Hippo pathway activates Dronc, which induces non-cell-autonomous mTOR activation not through Drice/Dcp1. Dronc regulates not only Drice/Dcp1 for apoptosis but also Rho1-Wingless (Wg)/Spitz signaling for compensatory proliferation, a phenomenon in which the dying cells promote cell proliferation in the live surrounding cells (Fan et al. 2014). To investigate the involvement of Rho1-Wg/Spitz signaling, we knocked down *rho1*, *wg*, or *spitz* in the Hippo-activated cells. As a result, we found that the non-cell-autonomous mTOR activation was significantly suppressed (Fig. 4E, H–J, quantified in F, K). Moreover, Wg was expressed in different patterns and ectopically expressed by Hippo activation compared with control (Fig. 4L and M). In addition to the ectopic Wg expression tending to occur in the hinge/ventral notum but not the pouch region (Fig. 4L and M), Wg expression level significantly increased by Hippo activation in the hinge/ventral notum but not the pouch region (Fig. 4N–Q). These data indicate that the Hippo-activated cells would drive Dronc-Rho1 signaling to promote the expression of *wg*/*spitz* in especially the hinge/ventral notum region for non-cell-autonomous tumorigenesis.

**Figure 4.**
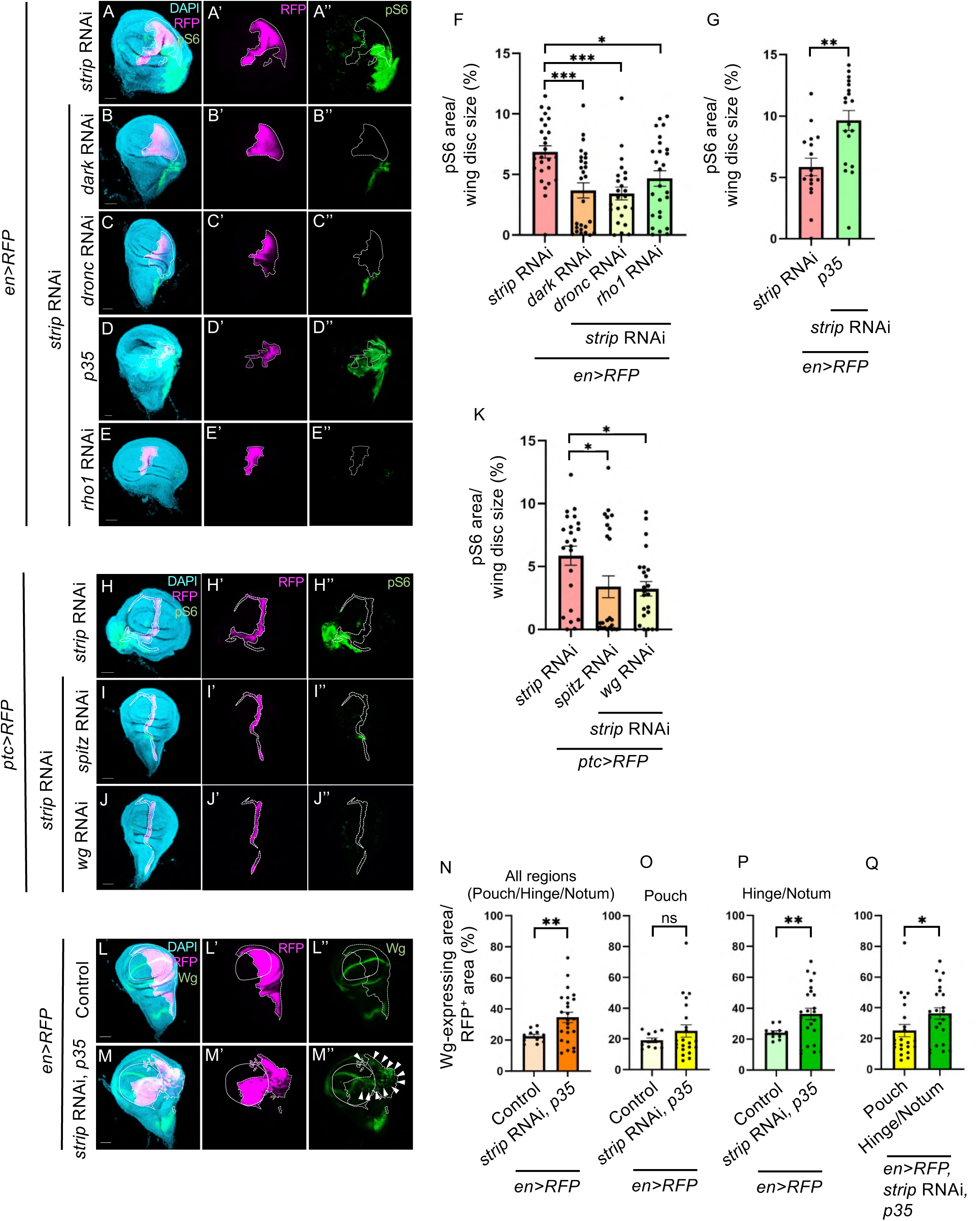
Hippo activation induces the mTOR-activated tumors through Dronc-Wg/Spitz signaling. (A–E) Confocal images show the wing discs bearing *strip-*knockdown cells with or without *dark*, *dronc*, *rho1* knockdown, or *p35* overexpression marked with RFP expression (magenta) and stained with anti-phospho-S6 (green). RFP (magenta) outlined by white dashed lines marks the expression pattern of *en-Gal4* in the wing disc. (F–G) Quantification of the size of the phospho-S6 positive region (% of phospho-S6 positive area/disc area) in the wing disc bearing wild-type, *strip-*knockdown cells with or without *dark*, *dronc*, *rho1* knockdown, or *p35* overexpression. ∗∗∗p < 0.001; ∗p < 0.05; one-way ANOVA with Dunnett’s multiple comparison test for (F). ∗∗p < 0.01; unpaired t-test for (G). (H–J) Confocal images show the wing discs bearing *strip-*knockdown cells with or without *spitz* or *wg* knockdown marked with RFP expression (magenta) and stained with anti-phospho-S6 (green). RFP (magenta) outlined by white dashed lines marks the expression pattern of *ptc-Gal4* in the wing disc. (K) Quantification of the size of the phospho-S6 positive region (% of phospho-S6 positive area/disc area) in the wing disc bearing wild-type, *strip-*knockdown cells with or without *wg* or *spitz* knockdown. ∗p < 0.05; one-way ANOVA with Dunnett’s multiple comparison test. (L–M) Confocal images show the wing discs bearing wild-type or *strip-*knockdown cells with *p35* overexpression marked with RFP expression (magenta) and stained with anti-Wg (green). RFP (magenta) outlined by white dashed lines marks the expression pattern of *en-Gal4* in the wing disc. Solid lines outline the pouch region. White arrowheads indicate ectopic expression of Wg. (N–Q) Quantification of the Wg-expressing area (% of Wg-expressing area/RFP-positive area) in the wing disc bearing wild-type or *strip-*knockdown cells with p35 overexpression. Since *en-Gal4* driver was active in the pouch, hinge, and ventral notum, the RFP-positive area could be separated into two areas, the pouch and hinge/ventral notum region. Quantification was conducted in the RFP-positive area of all regions (the pouch/hinge/ventral notum region), the pouch, or the hinge/ventral notum region. ∗∗p < 0.01; Welch’s t-test for (N–P). ∗p < 0.05; Unpaired t-test for (Q). Scale bars represent 50 μm.

### The activation of Dronc-Wg/Spitz signaling alone is insufficient to induce non-cell-autonomous tumorigenesis

Given that Dronc-Wg/Spitz signaling is used in compensatory proliferation, we examined whether the non-cell-autonomous tumorigenesis by the Hippo-activated cells is simply caused by compensatory proliferation. Thus, we investigated whether the activation of the key signaling for compensatory proliferation, Dronc-Wg/Spitz signaling, alone is sufficient to induce non-cell-autonomous tumorigenesis. To this end, we expressed *hid* or *rpr* with *p35*, which can activate Dronc-Wg/Spitz signaling without apoptosis (Ryoo, Gorenc, and Steller 2004; Fan et al. 2014, Fig. 5A). These cells could unexpectedly induce cell-autonomous mTOR activation, but they could not significantly induce non-cell-autonomous mTOR activation, as compared to that shown by the Hippo-activated cells (Fig. 5C–F, quantified in G). Moreover, even in the vicinity of cells ectopically expressing Wg, non-cell-autonomous mTOR activation was not significantly induced (Fig. 5H and I). These data suggest that non-cell-autonomous tumorigenesis by the Hippo-activated cells is not simply caused by compensatory proliferation. This observation suggests that Wg expression alone would be insufficient to activate the mTOR pathway non-cell-autonomously. Given the additional factors, including *atg8a*, *cyclin E*, and *bantam*, were required for Hippo activation-mediated tumorigenesis, there may be different secretory molecules in addition to Wg/Spitz (Fig. 5B)

**Figure 5.**
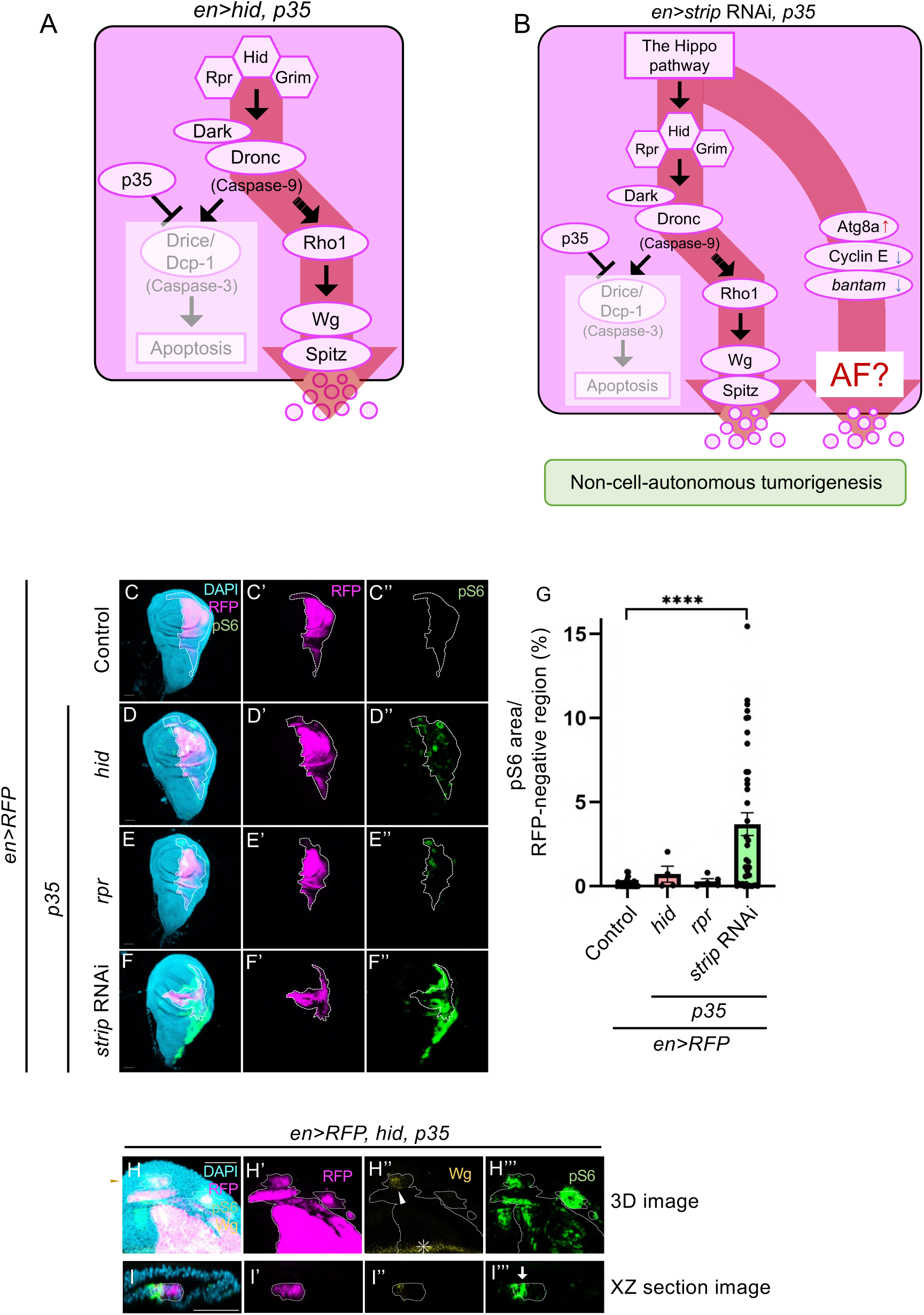
Activation of Dronc-Wg/Spitz signaling alone could not significantly induce non-cell-autonomous mTOR activation. (A) The schematic depiction indicates a driven signaling pathway in *hid*-overexpressing cells with *p35* overexpression. (B) The schematic depiction indicates a driven signaling pathway in *strip*-knockdown cells with *p35* overexpression. AF: additional factors. (C–F) Confocal images show the wing discs bearing wild-type, *strip-*knockdown, *hid*-, or *rpr*-overexpressing cells with *p35* overexpression marked with RFP expression (magenta) and stained with anti-phospho-S6 (green). RFP (magenta) outlined by white dashed lines marks the expression pattern of *en-Gal4* in the wing disc. (G) Quantification of the size of the phospho-S6 positive region (% of phospho-S6 positive area/RFP-positive area) in RFP-positive region bearing wild-type, *strip-*knockdown, *hid*, or *rpr* overexpressed cells with *p35* overexpression. ∗∗∗∗p < 0.0001; one-way ANOVA with Dunnett’s multiple comparison test. (H–I) A high-magnification confocal image shows the wing discs bearing *hid*-overexpressing cells with *p35* overexpression marked with RFP expression (magenta) and stained with anti-phospho-S6 (green) and anti-Wg (yellow). RFP (magenta) outlined by white dashed lines marks the expression pattern of *en-Gal4* in the wing disc. The brown arrowhead in the high magnification image indicates the cut position for the XZ section images. The white arrowhead indicates ectopic Wg expression. The white arrow indicates the cell-autonomous mTOR-activated cells. Scale bars represent 50 μm.

### Amino acid transporters Sat1/2 and Spitz redundantly act to achieve non-cell-autonomous tumorigenesis

Although the knockdown of *spitz* or *wg* using *the ptc-Gal4* driver significantly reduced the size of mTOR-activated tumors, the effects were limited (Fig. 4H–J, quantified in K). Moreover, the knockdown of *spitz* using the *en-Gal4* driver could not significantly reduce the size of the mTOR-activated tumors (Fig. 6I, quantified in N). Therefore, we assumed that there might be additional factors secreted from the Hippo-activated cells in addition to Wg/Spitz (Fig. 5B). Thus, we investigated the additional factors involved in Hippo activation-mediated tumorigenesis. Here, we conducted a genetic modifier screen with genomic deficiencies to identify the factors that genetically interact with the Hippo pathway. For this screening, we used the pupal lethality induced by the activation of the Hippo pathway (*strip* RNAi using *OK6-Gal4* driver that activates the expression in motor neurons) (Supplementary Fig. 6A). We screened the genes whose loss can rescue pupal lethality caused by Hippo activation and discovered the uncharacterized putative amino acid transporter, *CG4991*, which we named Sat1 (Supplementary Fig. 6A, quantified in B). This rescue rate increased when they are cultured in food with a high yeast concentration, suggesting that the level of protein/amino acid influences the Hippo activation-mediated pupal lethality. Interestingly, inside the *sat1* gene region, another gene encoding the uncharacterized putative amino acid transporter, *CG16700*, which we named *sat2*, was identified (Supplementary Fig. 6C). *Drosophila sat1/2* are conserved as *SLC36A1-4* in humans. Given that the *sat1* and *sat2* genes share the 5′ UTR and have similar amino acid sequences (Supplementary Fig. 6C and D), they are likely to work redundantly. As expected, the double knockdown of *sat1* and *sat2* more strongly suppressed the mTOR activation as compared to the single knockdown of *sat1* or *sat2* (Fig. 6A–E, quantified in F). These results indicate that *sat1* and *sat2* act redundantly in the Hippo-activated cells for non-cell autonomous tumorigenesis. Given that the amino acids activate the mTOR pathway (Laplante and Sabatini 2012), the Hippo-activated cells may secrete amino acids through Sat1/2 to activate the mTOR pathway in their surrounding cells. Since *sat1* was mainly localized in the plasma membrane and partially localized in the intracellular region in *sat1*-overexpressed S2 cells (Supplementary Fig. 7A-M), Sat1 in the plasma membrane has the potential to regulate amino acid levels in the extracellular region. To examine if the amino acids regulated by Sat1/2 could contribute to the non-cell autonomous tumorigenesis, we generated the double mutants for *sat1* and *sat2* (*sat1/2*) (Supplementary Fig. 6C) and conducted *spitz* knockdown in the *sat1/2* mutant background. Although *sat1/2* mutation or *spitz* knockdown alone partially or hardly reduced the frequency of non-cell-autonomous tumorigenesis (Fig. 6G–K, quantified in N), the combination of *sat1/2* mutation and *spitz* knockdown significantly suppressed the mTOR activation (Fig. 6L, M, quantified in N). This suggests that Spitz would work with Sat1/2 redundantly for Hippo activation-mediated tumorigenesis. Taken together, the Hippo-activated cells have the potential to secrete amino acids and Spitz/Wg to activate the mTOR pathway in their surrounding cells.

**Figure 6.**
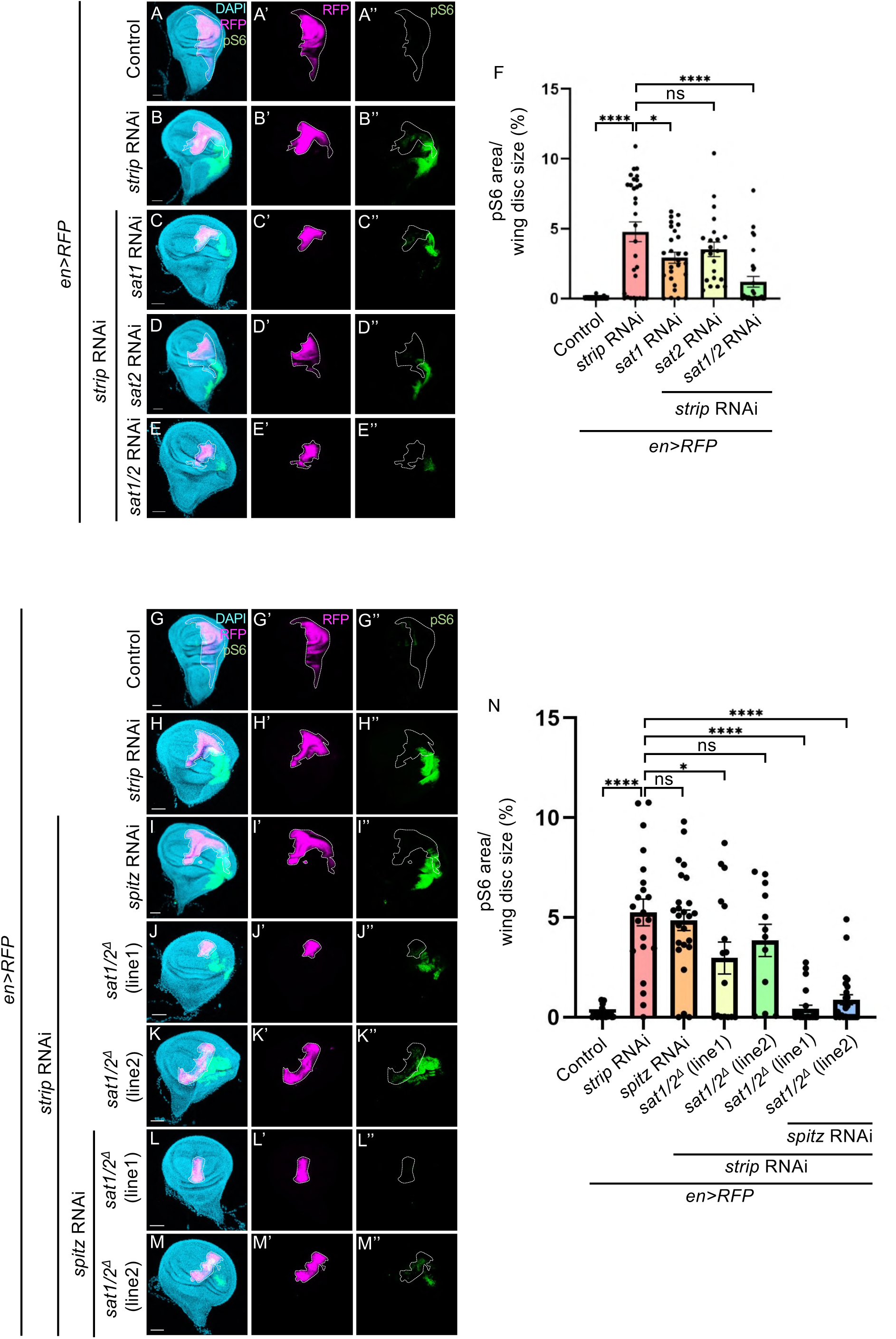
Hippo activation induces the mTOR-activated tumors through the amino acid transporter *sat1/2*. (A–E) Confocal images show the wing discs bearing wild-type or *strip-*knockdown cells with or without *sat1*, *sat2*, or *sat1/2* knockdown marked with RFP expression (magenta) and stained with anti-phospho-S6 (green). RFP (magenta) outlined by white dashed lines marks the expression pattern of *en-Gal4* in the wing disc. (F) Quantification of the size of the phospho-S6 positive region (% of phospho-S6 positive area/disc area) in the wing disc bearing wild-type or *strip-*knockdown cells with or without *sat1*, *sat12*, or *sat1/2* knockdown. ∗∗∗∗p < 0.0001; ∗p < 0.05; one-way ANOVA with Dunnett’s multiple comparison test. (G–M) Confocal images show the wing discs bearing wild-type or *strip-*knockdown cells with or without *spitz* knockdown alone or *spitz* knockdown with the *sat1/2* background mutation marked with RFP expression (magenta) and stained with anti-phospho-S6 (green). The male animals of these strains were selected for the experiment and the sat1/2 background mutation was hemizygous. RFP (magenta) outlined by white dashed lines marks the expression pattern of *en-Gal4* in the wing disc. (N) Quantification of the size of the phospho-S6 positive region (% of phospho-S6 positive area/disc area) in the wing disc bearing wild-type, *strip-*knocked down cells with or without *spitz* knockdown alone or *spitz* knockdown with the *sat1/2* background mutation. ∗∗∗∗p < 0.0001; ∗p < 0.05; one-way ANOVA with Dunnett’s multiple comparison test. Scale bars represent 50 μm.

## Discussion

Many studies have asserted that the Hippo pathway acts as a tumor suppressor in *Drosophila* and mammals because the mutations in the Hippo pathway promote tumorigenesis (Moroishi, Hansen, and Guan 2015; Yu, Zhao, and Guan 2015). However, recent studies have suggested that the Hippo pathway acts as a tumor promoter. For example, the Hippo pathway promotes tumor growth and invasive behavior in a cell-autonomous manner (Moroishi et al. 2016; Cottini et al. 2014; Yuan et al. 2008; Pearson et al. 2021). Here, we provide genetic evidence indicating that the Hippo-activated cells act as oncogenic niche cells, which promote tumorigenesis in a non-cell-autonomous manner (Fig. 7).

**Figure 7.**
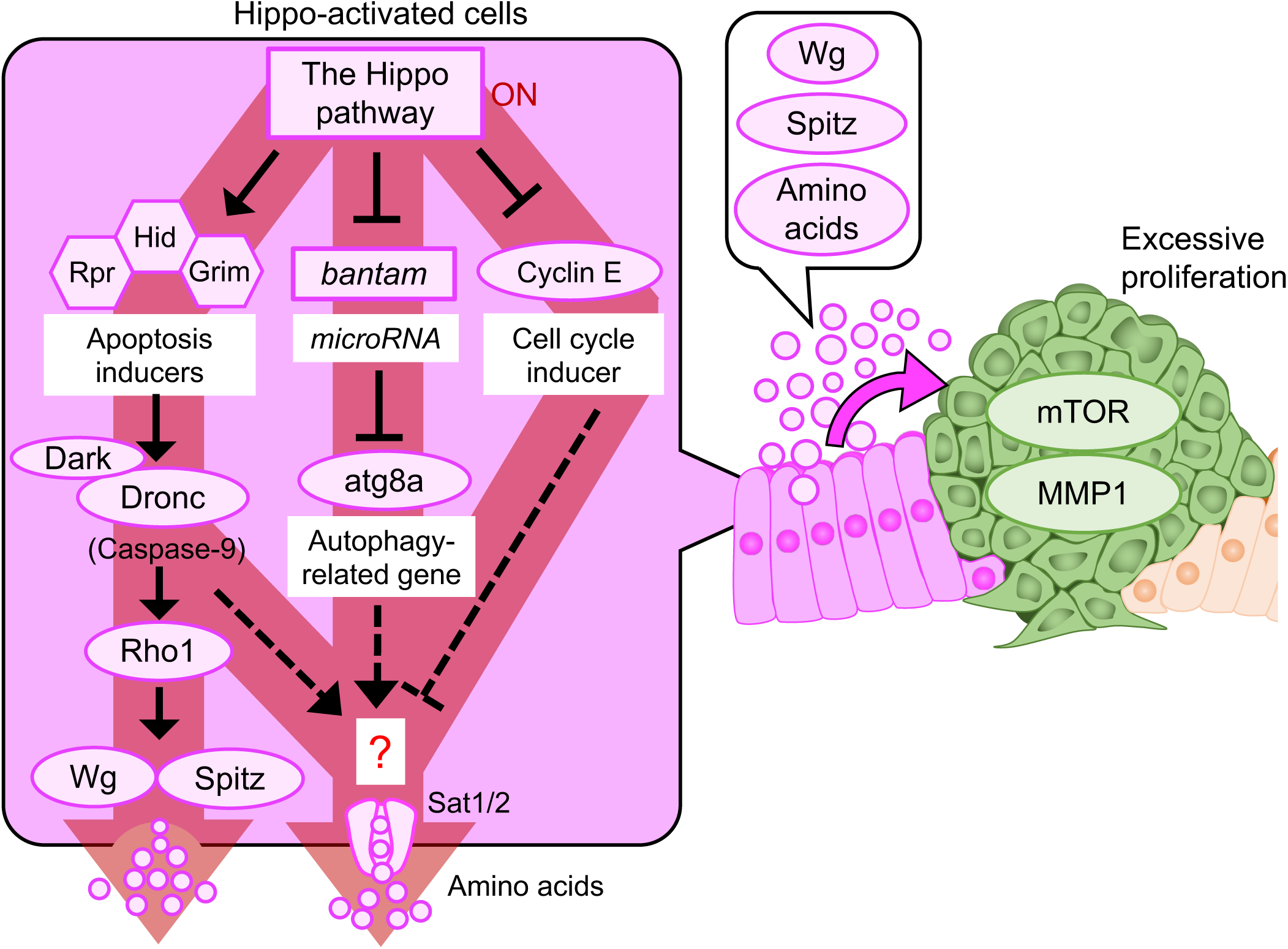
Hippo-activated cells induce non-cell-autonomous tumorigenesis through Spitz, Wg, and Sat1/2. The schematic depiction is the mechanistic hypothesis underlying the non-cell-autonomous tumorigenesis caused by the Hippo-activated cells. Hippo activation changes several gene expressions/activities, including *hid*, *bantam*, *atg8a*, and *cyclin E*. These alterations promote non-cell-autonomous tumorigenesis through Wg/Spitz secretion and Sat1/2.

Previous studies have suggested that tumor heterogeneity is crucial for tumor progression: cell-cell communication among oncogenic cells, including Src-activated and Ras-mutated cells, promotes invasive behaviors (Enomoto, Takemoto, and Igaki 2021). Given that Hippo-activated cells are crucial for tumorigenesis through cell-cell communication (Fig. 7), Hippo-activated cells may also heterogeneously exist in human tumors. To check this hypothesis: whether there are heterogeneously Hippo-activated cells in human tumors, we used spatial transcriptome data of colorectal tumors. We found that the Hippo pathway is likely activated heterogeneously in human colorectal tumors (Supplementary Fig. 8A–D, analyzed with data from F. Wang et al. 2023), suggesting that the Hippo-activated cells may communicate with their surrounding cells (non-Hippo-activated cells) for tumor progression.

Previous studies have found that the Src-activated cells act as oncogenic niche cells that induce non-cell-autonomous tumorigenesis through the activation of the JNK pathway (Enomoto and Igaki 2013; Enomoto, Vaughen, and Igaki 2015). Given that Rho1, a regulator of the JNK pathway, was required in the Hippo-activated cells for non-cell-autonomous tumorigenesis (Fig. 4E, quantified in F), a common mechanism might be present between the Hippo- and Src-activated cells. Moreover, the Src activation, which cooperates with *Ras^V12^* expression, promotes Wg expression through the JNK pathway (Hirabayashi, Baranski, and Cagan 2013). Given that the Hippo-activated cells upregulate Wg expression for non-cell-autonomous tumorigenesis (Fig. 4M, quantified in N–Q), the Wg expression might also be a common mechanism between the Hippo- and Src-activated cells. However, the Src activation inhibits the Hippo pathway (Enomoto and Igaki 2013), indicating that the Hippo pathway activity is different and opposite between the Hippo- and Src-activated oncogenic niche cells. Thus, these oncogenic niche cells would use the other pathways to promote non-cell-autonomous tumorigenesis.

We found that the region where Hippo is activated is important for tumor induction: the mTOR-activated tumors were induced by the Hippo activation in the hinge/ventral notum but not the pouch region (Fig. 2). We assume the reason why tumorigenesis by Hippo activation is more likely to occur in the hinge/ventral notum would be that Wg can be induced by Hippo activation there. Interestingly, Hippo activation seemed to drive Dronc activation in the pouch region, as shown by activated Drice/Dcp1 (Supplementary Fig. 3 G, H, quantified in I, J). Thus, there would be a mechanism in the pouch region that prevents activated Dronc from inducing Wg expression when the Hippo pathway activates Dronc.

The activation of Dronc-Wg/Spitz signaling alone did not exhibit Hippo activation-mediated tumorigenesis. Thus, we predicted that the Hippo-activated cells would use additional secretory molecules in addition to Wg/Spitz. We identified Sat1/2, a putative amino acid transporter, as the candidate regulators of additional secretory molecules. We believe that the Hippo-activated cells secrete amino acids through Sat1/2, which redundantly work with Wg/Spitz. Moreover, we found that the autophagy-related gene *atg8a* was crucial for tumorigenesis caused by Hippo activation. Given that autophagy activation promotes amino acid secretion, which promotes tumor progression in the surrounding benign tumors (Katheder et al. 2017; Sousa et al. 2016), the Hippo-activated cells may secrete amino acids through autophagy activation. Thus, the amino acids would be the leading candidates of secretory molecules for non-cell-autonomous mTOR activation.

The present study revealed the tumor-promoting role of the Hippo pathway, which is an unanticipated aspect of this pathway. This understanding will be useful in developing cancer treatments targeting the Hippo pathway.

## Materials and methods

### Fly strains

The flies were raised on standard fly food at 25°C. At 48–72 h after laying eggs, the heat shock for *hs-FLP*; *Ay-Gal4* was conducted for 30 min at 37°C. Unless otherwise stated, the sex of the larvae dissected was not differentiated. The following flies were used in this study: *ptc-Gal4* (BL2017), *en-Gal4*, *UAS-RFP* (BL30557), *ci-Gal4*, *UAS-GFP* (from Shizue Ohsawa), *pnr-Gal4* (BL3039), *nub-Gal4* (BL86108), *41D11-Gal4* (Medina, Calleja, and Morata 2021), *bx-Gal4* (BL8696), *284-Gal4* (Medina, Calleja, and Morata 2021), *OK6-Gal4* (Aberle et al. 2002), *hs-FLP*; *Ay-Gal4* (from Tatsushi Igaki), *UAS-strip* RNAi^#6+9^ (Sakuma et al. 2016), *UAS-strip* RNAi^#9-5^ (Sakuma et al. 2016), *UAS-wts* (Sansores-Garcia et al. 2011), *UAS-yki* RNAi (BL34067), *UAS-hippo* RNAi (BL33614), *UAS-bantam* (BL60671), *UAS-cyclin E* (BL4781), *UAS-atg8a* RNAi (BL34340), *UAS-miRHG* (From Chun-Hong Chen), *UAS-myc* (BL64759), *UAS-brk* (BL93081), *UAS-chinmo* RNAi (BL26777), *UAS-fng* RNAi (BL25947), *UAS-diap1* (BL63820), *UAS-InR*^#1^ (BL8248), *UAS-InR*^#2^ (BL8250), *UAS-dark* RNAi (BL33924), *UAS-dronc* RNAi (BL32963), *UAS-rho1* RNAi (BL27727), *UAS-p35* (BL5073), *UAS-p35* (from Bruce Hay), *UAS-hid* (BL65403), *UAS-rpr* (BL5824), *UAS-spitz* RNAi (BL28387), *UAS-wg* RNAi (BL31249), *UAS-CG4991* RNAi (VDRC30264), *UAS-CG16700* RNAi (BL61217), *sgRNA* for *sat2* (BL82722), nos-Cas9 (CAS-0001), *w; Sco/CyO, tubP-Gal80* (BL9491), UAS-mCD8-RFP (BL32218), UAS-mCD8-GFP (from Liqun Luo), and *Drosophila* deficiency lines (Bloomington *Drosophila* Stock Center and Kyoto *Drosophila* Stock Center).

The *sat1/2* mutant was generated in this study. As shown in Supplementary Fig. 6C, the *sat1* gene region was replaced by *3 × P3 DsRed* to generate a *sat1* null mutant using CRISPR-Cas9-triggered homologous recombination. Two guide RNA vectors and a donor vector were injected in the *vasa-Cas9* fly line (GenetiVision). The DNA fragments for guide RNAs (5′-CTTCGTACTTCATCGCTACTTCTC–3’, 5’- CTTCGCCGTGCTGGGCATCGTCAC-3′) were subcloned into the *Bbs*I-digested U6b-*sgRNA* vector. The 5’ homology arm (1 kbp-sequences upstream from 78 bp after the start codon) was amplified by PCR using the following primers: 5′-CCCTTCGCTGAAGCAGGTGGctcacagcgagcaagaggtgatcatcg-3′ and 5′-GCAGGTGTGCATATGTCCGCaagtagcgatgaagtacagttaggtctc-3′. The 3′ homology arm (1 kbp-sequences downstream from 46 bp before the stop codon) was amplified by PCR using the following primers: 5′-TATAGAAGAGCACTAGTAAAcactggcacctaccagagcatcgtgg-3′ and 5′-ACTCGATTGACGGAAGAGCCtactttttaaaaaacccctattaccc-3′. The 5’ homology arm was subcloned into the cut site of the *pHD-DsRed-attP* vector, which was cut using the *EcoR*L and *Not*I. The 3’ homology arm was subcloned into the cut site of the *pHD-DsRed-attP* vector, which *Bgl*II and *Xho*L. We screened the DsRed-positive transformants, and the *sat1* depletion was confirmed by genomic sequencing. Then, we crossed the *sat2 sgRNA* line (BL82722) with flies expressing *Cas9* in the germ cells (CAS-0001) to induce *sat2* mutations in the *sat1* mutant background. In the F1 generation, a heteroduplex mobility assay was conducted using microchip electrophoresis on MultiNA (Shimazu, MCE-202). Finally, we confirmed the *sat2* mutations by genomic sequencing.

### Immunostaining

The wandering third-instar larvae were dissected in phosphate-buffered saline (PBS) containing 0.3% Triton X-100 (0.3% PBST), fixed with 4% paraformaldehyde (PFA) for 20 min at room temperature, and washed three times with 0.3% PBST for 20 min. Then, these samples were blocked with 5% Normal Goat Serum (NGS) in 0.3% PBST for 1 h. The blocked samples were incubated at 4°C overnight with primary antibodies in 0.3% PBST, washed three times with 0.3% PBST for 20 min, and then incubated at 4°C overnight with secondary antibodies in 0.3% PBST. After washing three times with 0.3% PBST, the samples were incubated at 4°C overnight or room temperature for 1 hour with a VECTASHIELD Mounting Medium (Funakoshi Corporation #H-1000) containing DAPI (1:100, Nacalai Tesque #11034-56, 0.15 μg/ml), and were mounted with same mounting medium. The primary antibodies used were rabbit anti-cleaved *Drosophila* Drice/Dcp-1 (Asp216) (1:100 or 1:400, Cell Signaling Technology #9578), rabbit anti-phospho-S6 (1:400, gifts from Jongkyeong Chung and Aurelio Teleman), mouse anti-MMP1 (1:1:1 mixture of 5H7B11, 3A6B4, and 3B8D12 were diluted 1:10, DSHB), mouse anti-α-tubulin (1:500, Cedarlane #CLT9002), rat anti-E-Cad (1:100, DSHB #DCAD2), mouse anti-Wg (1:100 DSHB #4D4), guinea pig anti-Expanded (1:5000, gifts from Rick Fehon), and mouse anti-Flag (1:200, Sigma-Aldrich #F1804-200UG). The secondary antibodies used were Goat anti-rabbit Alexa Fluor™ 568 (1:250 Invitrogen #A-11036), Goat anti-Rabbit Alexa 488 (1:250, Invitrogen #A-11034), Goat anti-Mouse Alexa 633 (1:250, Invitrogen #A-21052), Goat anti-Mouse Alexa 488 (1:250, Invitrogen #R37120), and Goat anti-guinea pig Alexa 488 (1:1000, Invitrogen #A-11073). Actin staining was conducted using TRITC-phalloidin (1:200, Sigma-Aldrich #P1951). The images were taken with a Zeiss LSM900 confocal microscope.

### EdU staining

We followed the protocol for Click-iT™ EdU Cell Proliferation Kit for Imaging, Alexa Fluor™ 647 dye (ThermoFisher #C10340). The wing discs were incubated in 1× PBS supplemented with EdU (1:1000) for 20 min at room temperature and washed one time with 1× PBS. The discs were fixed with 4% PFA for 30 min, washed three times with 0.3% PBST for 20 min, and blocked with 5% NGS in 0.3% PBST for 1 h at room temperature. To visualize EdU incorporation, the discs were incubated in the Click-iT^®^ reaction cocktail for 30 min and washed three times with 0.3% PBST for 20 min at room temperature. After EdU staining, the discs underwent antibody staining, following the protocol of immunostaining described above.

### Processing, calculation, and statistical analyses for image quantification

The images were processed by using the tool threshold to eliminate background by using ImageJ software. We defaulted the region of the wing disc, RFP^+^, or mTOR tumors as ROI (Region of Interest) using a tool of ImageJ software for analyzing particles. We measured the size of the whole wing disc, RFP^+^ region, RFP^-^ region, or mTOR tumors. We also measured the positive area of anti-phospho-S6, anti-MMP1, EdU, anti-Drice/Dcp1, anti-Expanded, or anti-Wg staining in the specific region (Whole wing disc, RFP^+^ region, RFP^-^ region, or mTOR tumors). The positive area of anti-phospho-S6, anti-MMP1, EdU, anti-Drice/Dcp1, anti-Expanded, or anti-Wg staining was divided by measured region (Whole wing disc, RFP^+^ region, RFP^-^ region, or mTOR tumors) for normalization. All statistical analyses were performed using GraphPad Prism 9 or 10. The quantitative data were statistically analyzed by Welch’s t-test, unpaired t-test for single comparison, or one-way analysis of variance (ANOVA) with Dunnett’s multiple comparison test for multiple comparisons. All experiments demonstrated in the same graph were conducted at the same time, and each experiment was conducted using more than 10 samples of wing discs.

### Genetic modifier screen to identify the strip-interacting genes

*Strip* knockdown in motor neurons using *OK6-Gal4* exhibits pupal lethality. We screened the genes whose deletion could rescue pupal lethality caused by *strip* knockdown. *UAS-strip RNAi; OK6-Gal4/ tubP-Gal80, CyO* was crossed with genomic deletion strains. In the F1 generation, the *strip* knockdown was conducted with the heterozygous genomic deletion background. We marked the pupae of the F1 generation and counted the number of adults that were enclosed by 120 h after pupation. The individuals that did not emerge within 120 h after pupation were determined as dead pupae. The deletion strains showing a survival rate of >10% were identified as strains that could rescue the pupal lethality caused by *strip* knockdown. For this screening, we used nutrient-rich food to make the growth of larvae easier. More details of food conditions will be sent upon request. The initial part of this screening was performed by Tomoki Umehara (University of Tokyo).

### Establishment of S2 cells bearing pMT-Flag-Sat1-puro and analysis of Sat1 localization

The Flag sequence was attached to the Sat1 sequence (DGRC#UFO02039), which was subcloned into *pMT-puro* (addgene#17923) plasmid. *pMT-Flag-Sat1-puro* plasmid or empty plasmid (*pMT-puro*) as control was transfected into S2 cells and inserted into the genome of S2 cells. These transfected S2 cells were selected by puromycin, and S2 cells bearing *pMT-Flag-Sat1-puro* or *pMT-puro* were established. After these S2 cells (2.0L×L10^6^ cells/well) were seeded on a 6-well plate, these cells were incubated with 0.2LmM CuSO_4_(II) for 24 h to activate copper-inducible promoter (*metallothionein* gene promoter) and to induce Flag-Sat1 overexpression. By staining the FLAG-tag, Sat1 localization was detected.

### Data analysis of the spatial transcriptomics

We used data from previously conducted spatial transcriptomics of colorectal tumors (F. Wang et al. 2023). Data analysis was conducted following the method described in a previous study (F. Wang et al. 2023), and we classified the samples into the paratumor and tumor groups. We extracted data from the tumor group and evaluated the Hippo pathway activity using the YAP-target gene signature of 22 genes as described previously (Kowalczyk et al. 2022). We evaluated the Hippo pathway activity by assessing the average expression of 22 YAP-target genes in the colorectal tumor group.

## Acknowledgments

We thank Chun-Hong Chen (National Health Research Institute), Tatsushi Igaki (Kyoto Univ), Shizue Ohsawa (Nagoya Univ), Bloomington Drosophila Stock Center, and the Kyoto Drosophila Stock Center for the fly stocks. We thank Jongkyeong Chung (Seoul National Univ) and Aurelio Teleman (German Cancer Research Center) for the anti-phosphorylated pS6 antibody. We also thank Tomoki Umehara for the assistance in the genetic modifier screen. We are grateful to Enago (https://www.enago.com) for the English language editing. This work was supported by the Nozomi H Foundation, NOVARTIS Foundation (Japan) for the Promotion of Science, and JSPS KAKENHI (grant numbers 21H02479 and 24K02062) to T.C., JSPS KAKENHI (grant number 20K15903) to M.O., and JSPS Research Fellows (grant number: 24KJ1732) and Takenaka Scholarship Foundation to D.H.

## Author contributions

D.H., M.O. and T.C. conceived this project. D.H. performed the experiments, except for the genetic modifier screen. C.S. and Tomoki Umehara carried out the genetic modifier screen. T.C. and M.O. supervised this project. T.A. and M.M. contributed intellectual support on cancer biology and cell death signaling. The manuscript was written by D.H., M.O., and T.C. with inputs from all authors.

