## Supplemental information for "Hippo-activated cells induce non-cell autonomous tumorigenesis in *Drosophila*"

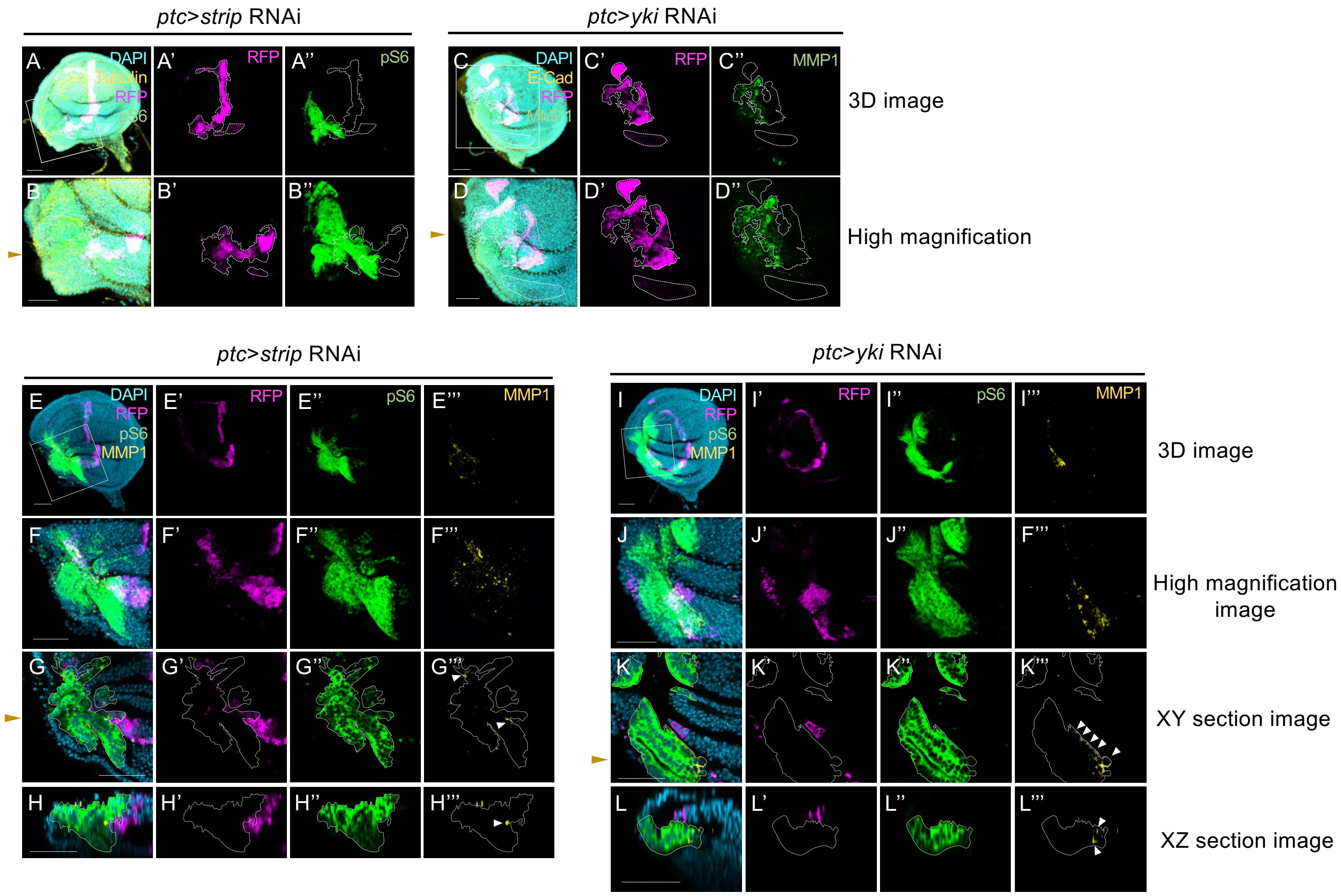


**Supplementary Fig. 1. MMP1 expression is induced in the mTOR-activated cells by Hippo activation**

(A and B) Confocal images show the wing discs bearing *strip*-knockdown cells marked with RFP expression (magenta) and stained with anti-phospho-S6 (green) and anti-tubulin (yellow). The 3D (A) and high magnification images (B) of the wing discs. The brown arrowhead in the high magnification image indicates the cut position for the XZ section images (Fig. 1F). RFP (magenta) outlined by white dashed lines marks the expression pattern of *ptc-Gal4* in the wing disc. Scale bars represent 50 µm. (C and D) Confocal images show the wing discs bearing *yki*-knockdown cells marked with RFP expression (magenta) and stained with anti-MMP1 (green) and anti-E-Cad (yellow). The 3D (C) and high magnification images (D) of the wing discs. The brown arrowhead in the XY section image (D) indicates the cut position for the XZ section images (Fig. 1 L). RFP (magenta) outlined by white dashed lines marks the expression pattern of *ptc-Gal4* in the wing disc. (E–L) Confocal images show the wing discs bearing *strip* or *yki*-knockdown cells marked with RFP expression (magenta) and stained with anti-phospho-S6 (green) and anti-MMP1 (yellow). The 3D (E and I), high magnification (F and J), XY section (G and K), and XZ section images (H and L) of the wing discs. The brown arrowhead in the XY section image indicates the cut position for the XZ section images. The white arrowhead shows MMP1-expressing cells in the mTOR-activated tumors outlined by white dashed lines. RFP (magenta) marks the expression pattern of *ptc-Gal4* in the wing disc. Scale bars represent 50 µm.

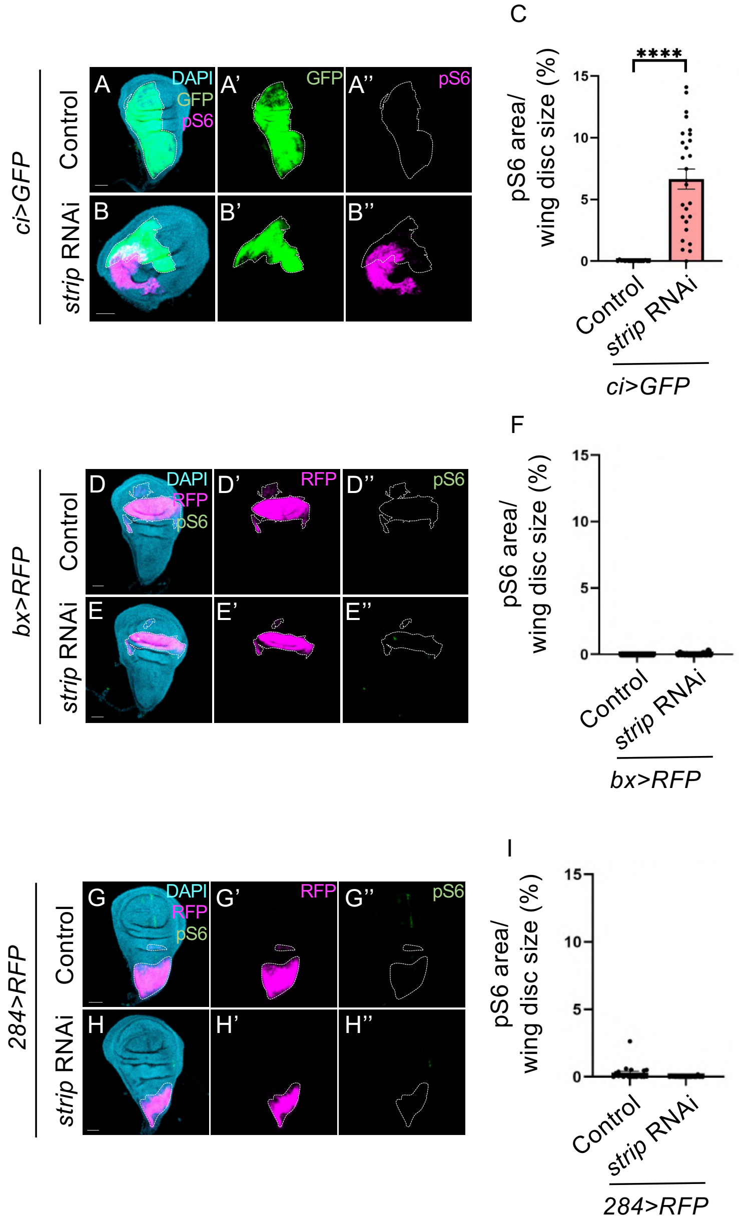


**Supplementary Fig. 2. Hippo activation in the pouch/dorsal notum region cannot induce the mTOR-activated tumors**

(A and B) Confocal images show the wing discs bearing wild-type or *strip*-knockdown cells marked with GFP expression (green) and stained with anti-phospho-S6 (magenta). GFP (green) outlined by white dashed lines marks the expression pattern of *ci-Gal4* in the wing disc. (D, E, G, H) Confocal images show the wing discs bearing wild-type or *strip*-knockdown cells marked with RFP expression (magenta) and stained with anti-phospho-S6 (green). RFP (magenta) outlined by white dashed lines marks the expression pattern of *bx-Gal4* or *284-Gal4* in the wing disc. (C, F, I) Quantification of the size of the phospho-S6 positive region (% of phospho-S6 positive area/disc area) in the wild-type or strip-knocked down wing disc. ****$p < 0.0001$; Welch's t-test. Scale bars represent 50 μm.

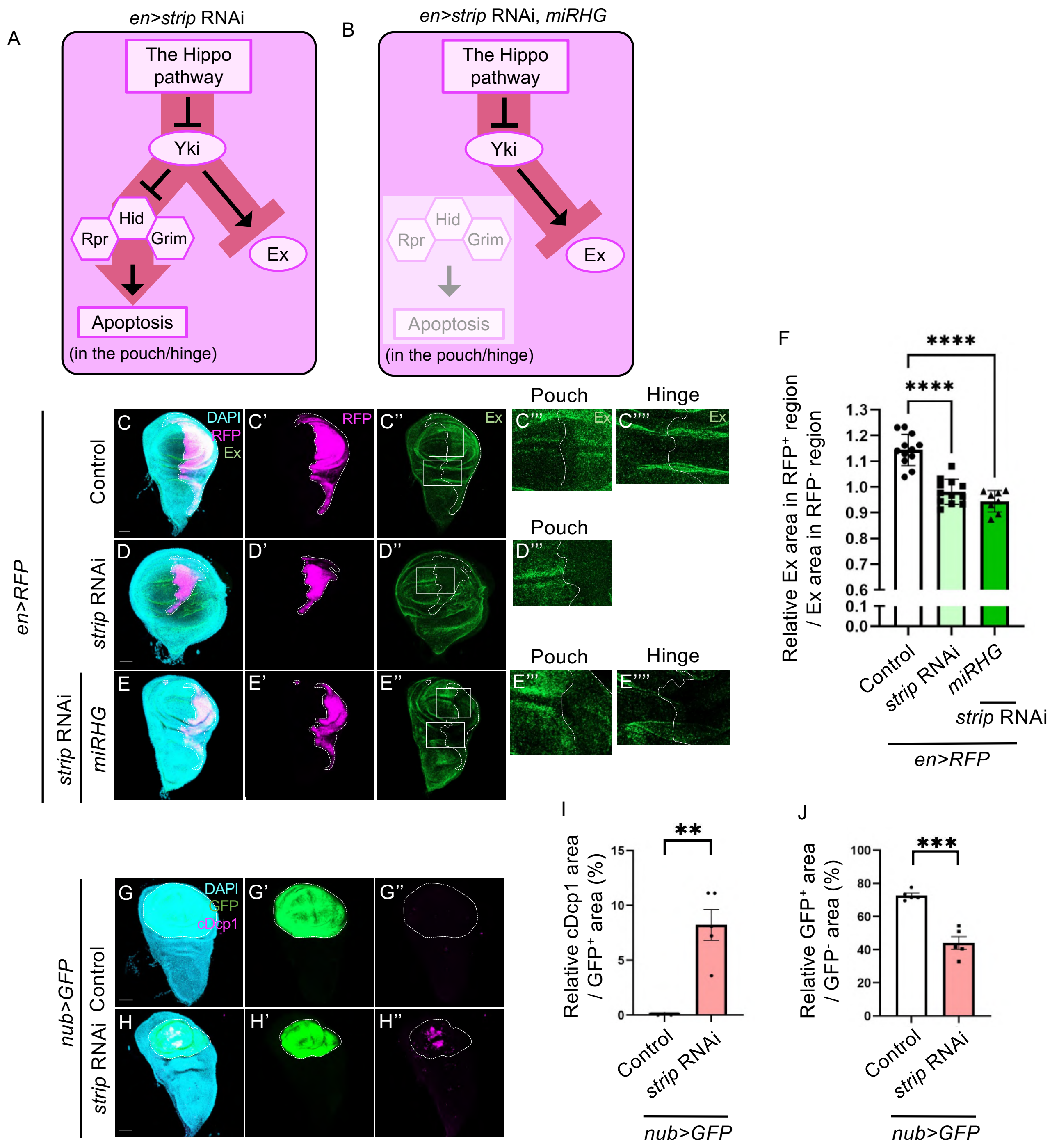


**Supplementary Fig. 3. The Hippo pathway is activated by *strip* RNAi in the pouch/hinge region**

(A and B) The schematic depiction indicates a driven signaling pathway in Hippo activation by *strip* RNAi with or without knockdown of *rpr*, *hid*, and *grim* (*miRHG*). *miRHG* suppresses apoptosis in the pouch and hinge region, which makes it easier to analyze these regions under Hippo activation. (C–E) Confocal images show the wing discs bearing wild-type or *strip*-knocked down cells with or without *miRHG* marked with RFP expression (magenta) and stained with anti-Expanded (green). RFP (magenta) outlined by white dashed lines marks the expression pattern of *en-Gal4* in the wing disc. (F) Quantification of the relative Expanded level (Expanded positive area in $RFP^+$ region/Expanded positive area in $RFP^-$ region) in the wild-type or strip-knockdown wing disc. $****p < 0.0001$; one-way ANOVA with Dunnett's multiple comparison test. (G–H) Confocal images show the wing discs bearing wild-type or *strip*-knockdown cells marked with GFP expression (green) and stained with anti-cleaved Drice/Dcp1 (Red). GFP (green) outlined by white dashed lines marks the expression pattern of *nub-Gal4* in the wing disc. (I) Quantification of the relative cleaved Drice/Dcp1 area (% of cleaved Drice/Dcp1 area/$GFP^+$ area) in the wild-type or *strip*-knocked down wing disc. $**p < 0.01$; Welch's t-test. (J) Quantification of the relative pouch size (% of $GFP^+$ area/$GFP^-$ area) in the wild-type or *strip*-knocked down wing disc. $***p < 0.001$; Unpaired t-test. Scale bars represent 50 μm.

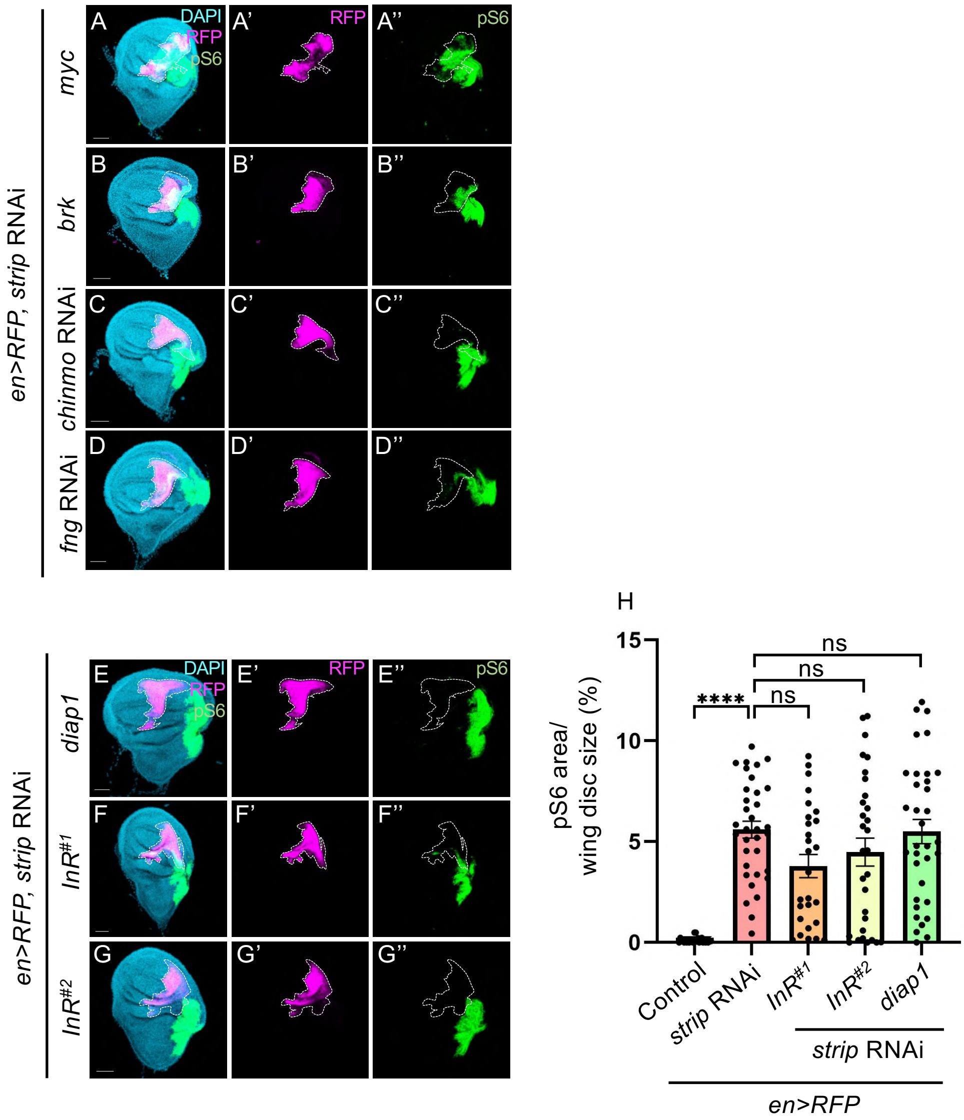


**Supplementary Fig. 4. Hippo activation, rather than by controlling the expression levels of *myc*, *brk*, *chinmo*, *fng*, *daip1*, and *InR,* induces the mTOR-activated tumors**

(A–G) Confocal images show the wing discs bearing wild-type or *strip*-knockdown cells with or without overexpression of *myc*, *brk*, *daip1*, *InR*, knockdown of *chinmo*, or *fng* marked with RFP expression (magenta) and stained with anti-phospho-S6 (green). RFP (magenta) outlined by white dashed lines marks the expression pattern of *en-Gal4* in the wing disc. (H) Quantification of the size of the phospho-S6 positive region (% of phospho-S6 positive area/disc area) in the wing disc bearing wild-type, *strip*-knockdown cells with or without overexpression of *daip1* or *InR*. ****p < 0.0001; one-way ANOVA with Dunnett's multiple comparison test. Scale bars represent 50 μm.

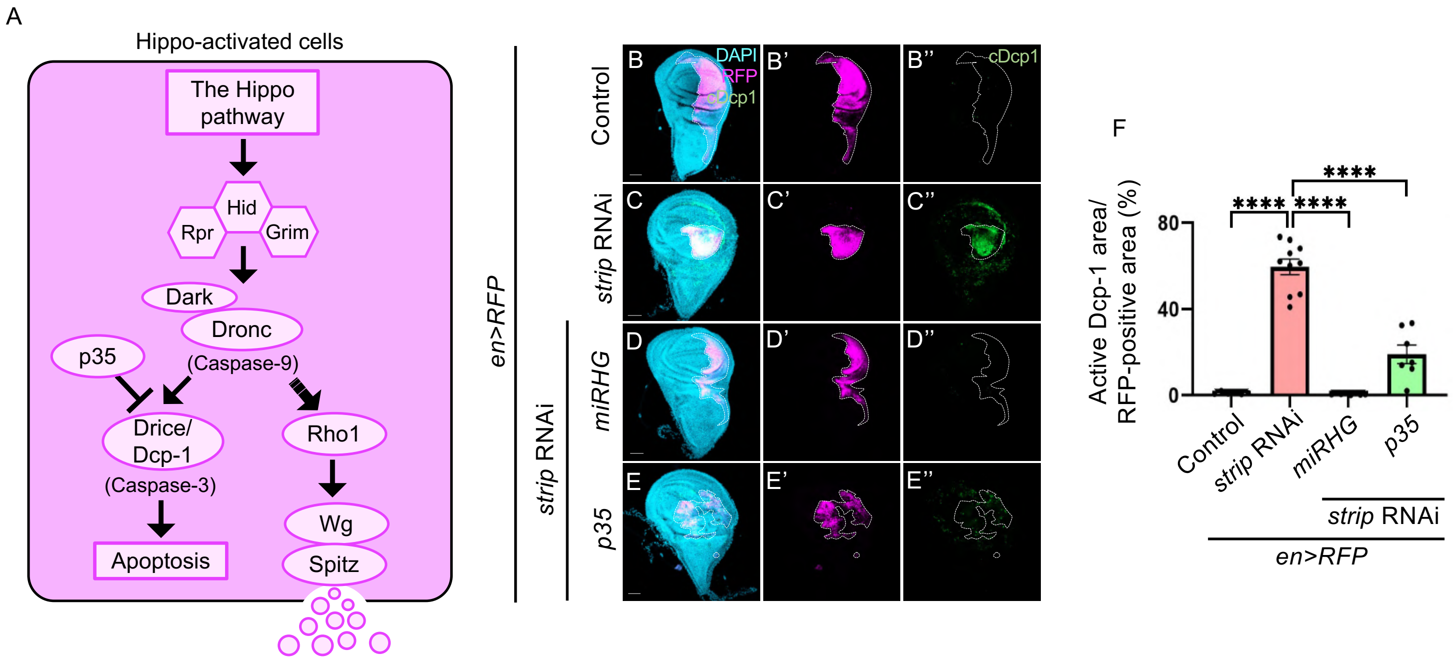


**Supplementary Fig. 5. Hippo activation by *strip* RNAi drives apoptosis signaling**

(A) The schematic depiction indicates the signaling pathway of apoptosis and compensatory proliferation. (B–E) Confocal images show the wing discs bearing wild-type or *strip*-knockdown cells with or without *rpr*, *hid*, and *grim* knockdown or *p35* overexpression marked with RFP expression (magenta) and stained with anti-cleaved Drice/Dcp1 (green). RFP (magenta) outlined by white dashed lines marks the expression pattern of *en-Gal4* in the wing disc. Knockdown of *rpr, hid,* and *grim* was conducted by the expression of microRNAs for *rpr*, *hid*, and *grim* (*miRHG*). (F) Quantification of the size of the cleaved Drice/Dcp1-positive region (% of anti-cleaved Drice/Dcp1 positive area/disc area) in the wing disc bearing wild-type or *strip*-knockdown cells with or without *pr*, *hid*, and *grim* knockdown or *p35* overexpression. ****p < 0.0001; one-way ANOVA with Dunnett's multiple comparison test. Scale bars represent 50 µm.

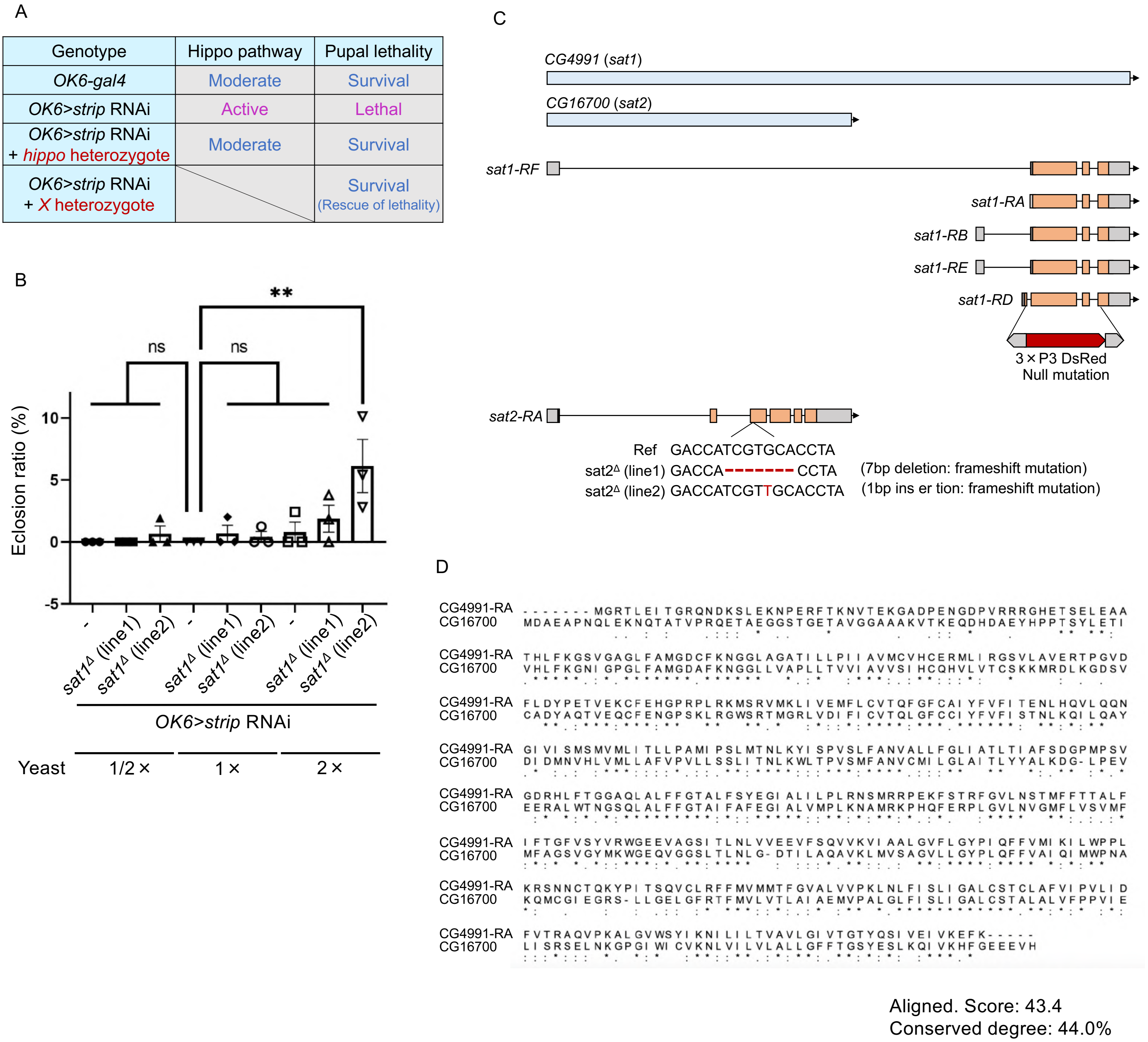


**Supplementary Fig. 6. Genomic deficiency screening identifies *Sat1/2***

(A) The table shows the Hippo activity and pupal lethality in individuals bearing wild-type and *strip*-knockdown cells with or without several gene background mutations. The knockdown of *strip* was conducted using the *OK6-Gal4* driver for genomic deficiency screening. *X* genes are putative genes whose deletion can rescue pupal lethality caused by Hippo activation. (B) Quantification of eclosion ratio (% of the number of adult flies/the number of pupae) in individuals bearing wild-type or *strip*-knockdown cells with or without *sat1/2* background mutation. **p < 0.01; one-way ANOVA with Dunnett's multiple comparison test. (C) The schematic depictions show the *CG4991* (*sat1*) and *CG16700* (*sat2*) genes and the mutated site of *sat1/2*. The coding region of the *sat1* gene was replaced by the marker gene 3xP3 DsRed. The coding region of the *sat2* gene was mutated with deletion (line 1) or insertion (line 2). (D) The similarity of the amino acid sequence between Sat1 and Sat2. Sat1 and Sat2 were aligned using ClustalW. Asterisk (*): sequence identity; colon (:): strongly similar properties; period (.): weakly similar properties.

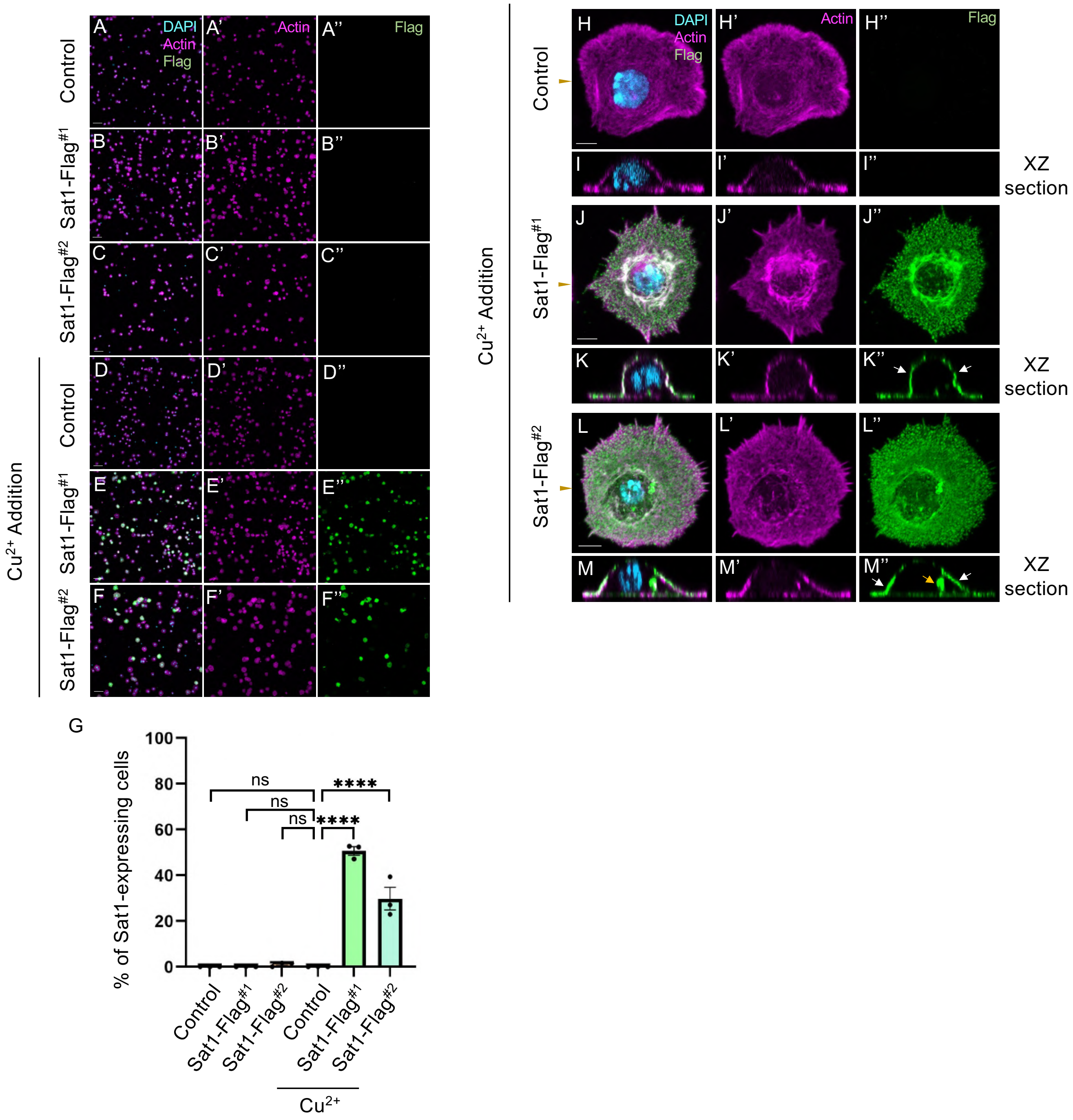


**Supplementary Fig. 7. Sat1 is mainly localized in the plasma membrane**

(A–F, H–M) Confocal images of low magnification (A–F) or high magnification (H–M) show S2 cells bearing *pMT-puro* (control) or *pMT-Flag-Sat1-puro*. $Cu^{2+}$ addition activated copper-inducible promoter and induced overexpression of Flag-tagged Sat1. Sat1-Flag$^{\#1}$ and Sat1-Flag$^{\#2}$ are S2 cells (line1) and (line2) bearing *pMT-Flag-Sat1-puro*. Outlines of cells are stained with TRITC-phalloidin. Sat1 expression was indicated by anti-Flag staining (green). The brown arrowhead in the 3D image (H, J, L) indicates the cut position for the XZ section images. The white arrow indicates Sat1 localization in the plasma membrane, and the yellow arrow indicates intracellular localization of Sat1. Scale bars represent 50 μm for (A–F) and 5 μm for (H–M). (G) Quantification of the ratio of Sat1-expressing cells (% of the number of Sat1-expressing cells/total number of cells). ****$p < 0.0001$; one-way ANOVA with Dunnett's multiple comparison test.

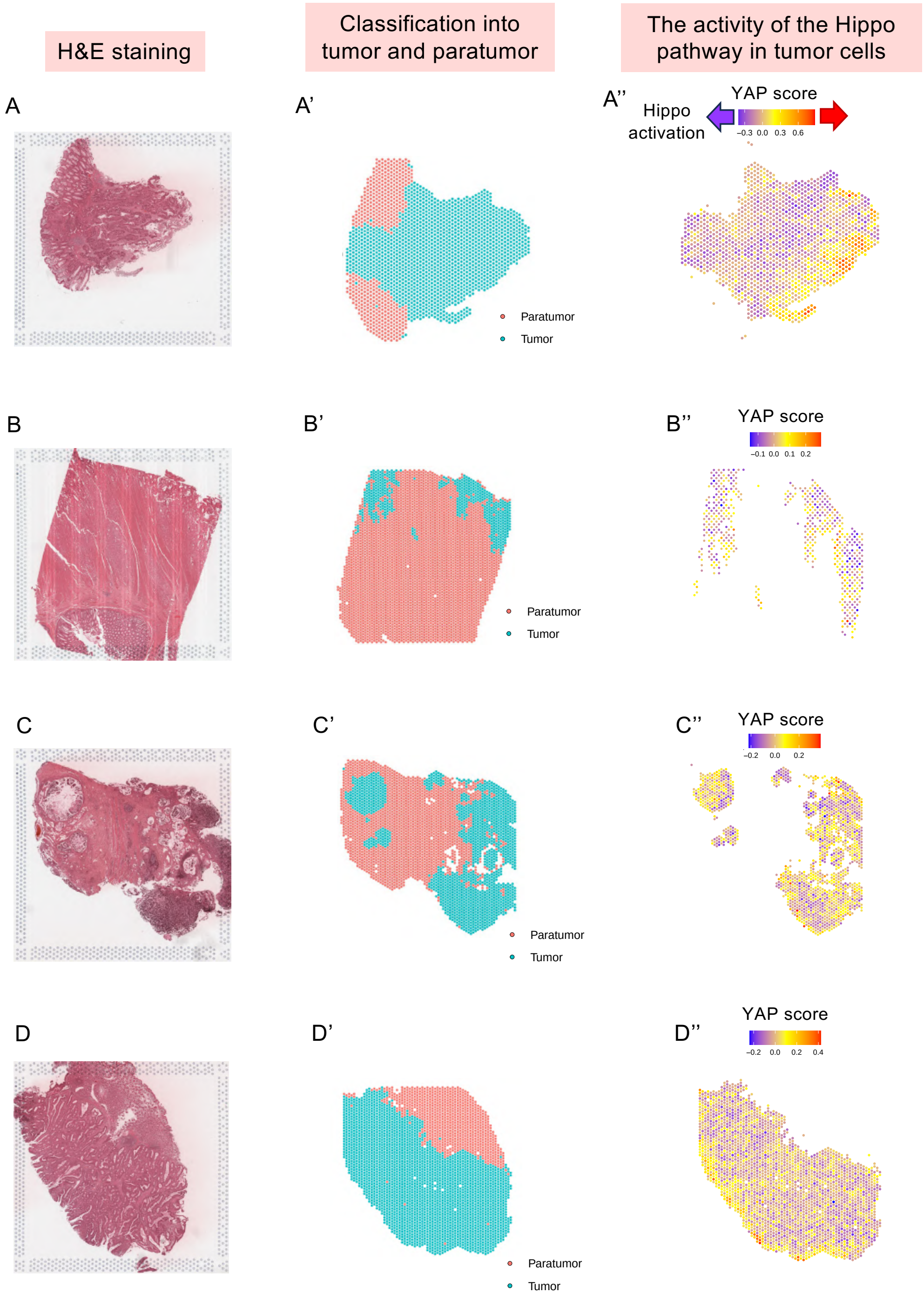


**Supplementary Fig. 8. Expression of YAP-target genes in the colorectal tumors**

Spatial transcriptome analysis was conducted using the data from the colorectal cancer group (F. Wang et al. 2023). (A–D) H&E staining shows a section of the colorectal cancer in each patient. (A'–D') Cells in the colorectal cancer section were classified into tumor or paratumor groups. (A''–D'') The activity of the Hippo pathway was evaluated by assessing the average expression of 22 genes of the YAP-target gene signature in the tumor cluster. The Hippo pathway is predicted to be active in tumors where the average expression of the YAP-target gene signature (YAP score) is low.

### Genotype list

Figure 1

(B, I) *ptc-Gal4/+; UAS-mCD8-RFP/ +*

(C,J, F) *ptc-Gal4/ UAS-strip RNAi; UAS-mCD8-RFP/ +*

(D) *ptc-Gal4/ UAS-wts; UAS-mCD8-RFP/ +*

(E, K, L) *ptc-Gal4/ +; UAS-mCD8-RFP/ UAS-yki RNAi*

(N) *en-Gal4, UAS-RFP/ +*

(O) *en-Gal4, UAS-RFP/ UAS-strip RNAi*

Figure 2

(B) *nub-Gal4/ +; UAS-mCD8-RFP/ +*

(C) *nub-Gal4/ UAS-strip RNAi ; UAS-mCD8-RFP/ +*

(D) *nub-Gal4/ +; UAS-mCD8-RFP/ UAS-yki RNAi*

(F) *pnr-Gal4/ UAS-mCD8-RFP*

(G) *UAS-strip RNAi/ + ; pnr-Gal4/ UAS-mCD8-RFP*

(I) *41D11-Gal4, UAS-GFP/ +*

(J) *UAS-strip RNAi/ +; 41D11-Gal4, UAS-GFP/ +*

(K) *41D11-Gal4, UAS-GFP/ UAS-yki RNAi*

(M, N) *hs-FLP/ + or y; Act>y>Gal4, UAS-GFP/ UAS-strip RNAi*

Figure 3

(B) *en-Gal4, UAS-RFP/ +*

(C) *en-Gal4, UAS-RFP/ UAS-strip RNAi*

(D) *en-Gal4, UAS-RFP/ UAS-strip RNAi; UAS-hippo RNAi/ +*

(E) *en-Gal4, UAS-RFP/ UAS-strip RNAi; UAS-bantam/ +*

(F) *en-Gal4, UAS-RFP/ UAS-strip RNAi; UAS-cyclinE/ +*

(G) *en-Gal4, UAS-RFP/ UAS-strip RNAi; UAS-atg8a RNAi/ +*

(H) *UAS-strip RNAi/ + or y; en-Gal4, UAS-RFP/ +*

(I) *UAS-strip RNAi/ + or y; en-Gal4, UAS-RFP/ UAS-miRHG*

Figure 4

(A) *en-Gal4, UAS-RFP/ UAS-strip RNAi*

(B) *en-Gal4, UAS-RFP/ UAS-strip RNAi; UAS-dark RNAi/ +*

(C) *en-Gal4, UAS-RFP/ UAS-strip RNAi; UAS-dronc RNAi/ +*

(D) *UAS-strip RNAi/ + or y ;en-Gal4, UAS-RFP/ UAS-p35*

(E) *en-Gal4, UAS-RFP/ UAS-strip RNAi; UAS-rho1 RNAi/ +*

(H) *ptc-Gal4/ UAS-strip RNAi; UAS-mCD8-RFP/ +*

(I) *ptc-Gal4/ UAS-strip RNAi; UAS-mCD8-RFP/ UAS-spitz RNAi*

(J) *ptc-Gal4/ UAS-strip RNAi; UAS-mCD8-RFP/ UAS-wg RNAi*

(L) *en-Gal4, UAS-RFP/ +*

(M) *UAS-strip RNAi/ + or y ;en-Gal4, UAS-RFP/ UAS-p35*

Figure 5

(C) *en-Gal4, UAS-RFP/ +*

(D, H, I) *en-Gal4, UAS-RFP/ UAS-hid; UAS-p35/ +*

(E) *en-Gal4, UAS-RFP/ UAS-rpr, UAS-p35*

(F) *UAS-strip RNAi/ + or y ;en-Gal4, UAS-RFP/ UAS-p35*

Figure 6

(A) *en-Gal4, UAS-RFP/ +*

(B) *en-Gal4, UAS-RFP/ UAS-strip RNAi*

(C) *en-Gal4, UAS-RFP/ UAS-strip RNAi; UAS-CG4991 RNAi/ +*

(D) *en-Gal4, UAS-RFP/ UAS-strip RNAi; UAS-CG16700 RNAi/ +*

(E) *en-Gal4, UAS-RFP/ UAS-strip RNAi; UAS-CG4991RNAi/ UAS-CG16700 RNAi*

(G) *en-Gal4, UAS-RFP/ +*

(H) *en-Gal4, UAS-RFP/ UAS-strip RNAi*

(I) *en-Gal4, UAS-RFP/ UAS-strip RNAi; UAS-spitz RNAi/ +*

(J) *sat1 Δ/2Δ (line1); en-Gal4, UAS-RFP/ UAS-strip RNAi*

(K) *sat1 Δ/2Δ (line2); en-Gal4, UAS-RFP/ UAS-strip RNAi*

(L) *sat1 Δ/2Δ (line1); en-Gal4, UAS-RFP/ UAS-strip RNAi; UAS-spitz RNAi/ +*

(M) *sat1 Δ/2Δ (line2); en-Gal4, UAS-RFP/ UAS-strip RNAi; UAS-spitz RNAi/ +*

Supplemetarry Fig. 1

(A, B, E–H) *ptc-Gal4/ UAS-strip RNAi; UAS-mCD8-RFP/ +*

(C, D, I–L) *ptc-Gal4/ +; UAS-mCD8-RFP/ UAS-yki RNAi*

Supplemetarry Fig. 2

(A) *ci-Gal4, UAS-GFP/ +*

(B) *ci-Gal4, UAS-GFP/ UAS-strip RNAi*

(D) *bx-Gal4/ + or y; ; UAS-mCD8-RFP/ +*

(E) *bx-Gal4/ + or y; UAS-strip RNAi/ +; UAS-mCD8-RFP/ +*

(G) *284-Gal4/ + or y; ; UAS-mCD8-RFP/ +*

(H) *284-Gal4/ + or y; UAS-strip RNAi/ +; UAS-mCD8-RFP/ +*

Supplemetarry Fig. 3

(C) *en-Gal4, UAS-RFP/ +*

(D) *UAS-strip RNAi/ + or y; en-Gal4, UAS-RFP/ +*

(E) *UAS-strip RNAi/ + or y; en-Gal4, UAS-RFP/ UAS-miRHG*

(G) *nub-Gal4/ UAS-mCD8-GFP*

(H) *UAS-strip RNAi; nub-Gal4/ UAS-mCD8-GFP*

Supplemetarry Fig. 4

(A) *en-Gal4, UAS-RFP/ UAS-strip RNAi; UAS-myc/ +*

(B) *en-Gal4, UAS-RFP/ UAS-strip RNAi; UAS-brk/ +*

(C) *en-Gal4, UAS-RFP/ UAS-strip RNAi; UAS-chimo RNAi/ +*

(D) *en-Gal4, UAS-RFP/ UAS-strip RNAi; UAS-fng RNAi/ +*

(E) *en-Gal4, UAS-RFP/ UAS-strip RNAi; UAS-diap1/ +*

(F) *en-Gal4, UAS-RFP/ UAS-strip RNAi; UAS-InR#1/ +*

(G) *en-Gal4, UAS-RFP/ UAS-strip RNAi; UAS-InR#2/ +*

Supplemetarry Fig. 5

(B) *en-Gal4, UAS-RFP/ +*

(C) *UAS-strip RNAi/ + or y; en-Gal4, UAS-RFP/ +*

(D) *UAS-strip RNAi/ + or y; en-Gal4, UAS-RFP/ UAS-miRHG*

(E) *UAS-strip RNAi/ + or y ;en-Gal4, UAS-RFP/ UAS-p35*

Supplemetarry Fig. 6

(B) *sat1 Δ (line1)/ + or y; OK6-Gal4/ UAS-strip RNAi*

(B) *sat1 Δ (line2)/ + or y; OK6-Gal4/ UAS-strip RNAi*

Supplemetarry Fig. 7

(A, D, H, I) *pMT-puro*

(B, E, J, K) *pMT-Flag-Sat1-puro (line1)*

(C, F, L, M) *pMT-Flag-Sat1-puro (line2)*
